# Development of antibiotic resistance reveals diverse evolutionary pathways to face the complex and dynamic environment of a long-term treated patient

**DOI:** 10.1101/2021.05.14.444257

**Authors:** Claudia A. Colque, Pablo E. Tomatis, Andrea G. Albarracín Orio, Gina Dotta, Diego M. Moreno, Laura G. Hedemann, Rachel A. Hickman, Lea M. Sommer, Sofía Feliziani, Alejandro J. Moyano, Robert A. Bonomo, Helle K. Johansen, Søren Molin, Alejandro J. Vila, Andrea M. Smania

**Affiliations:** Universidad Nacional de Córdoba, Facultad de Ciencias Químicas, Departamento de Química Biológica Ranwel Caputto, Córdoba, Argentina; CONICET, Universidad Nacional de Córdoba, Centro de Investigaciones en Química Biológica de Córdoba (CIQUIBIC), Córdoba, Argentina; Instituto de Biología Molecular y Celular de Rosario (IBR), CONICET, Universidad Nacional de Rosario, Rosario, Argentina; Area Biofísica, Facultad de Ciencias Bioquímicas y Farmacéuticas, Universidad Nacional de Rosario, Argentina; IRNASUS, Universidad Católica de Córdoba, CONICET, Facultad de Ciencias Agropecuarias, Córdoba, Argentina; IQUIR, Instituto de Química de Rosario, Universidad Nacional de Rosario, Santa Fe, Argentina; Department of Clinical Microbiology, Rigshospitalet, Copenhagen, Denmark; Novo Nordisk Foundation Centre for Biosustainability, Technical University of Denmark, Lyngby, Denmark; Department of Molecular Biology and Microbiology, Case Western Reserve University, Cleveland, Ohio, USA; Research Service, Louis Stokes Cleveland Department of Veterans Affairs, Cleveland, Ohio, USA; Department of Clinical Medicine, University of Copenhagen, Copenhagen, Denmark

**Keywords:** *Pseudomonas aeruginosa*, cystic fibrosis, β-lactamase evolution, ceftolozane resistance, hypermutability

## Abstract

Antibiotic resistance development has been studied using approaches that range from laboratory experimental evolution, surveillance and epidemiology, to clinical isolate sequencing. However, evolutionary trajectories depend on the environment in which selection takes place, compelling to address evolutionary analyses in antibiotic-treated patients, to embrace the whole inherent environmental complexities as well as their dynamics over time. Herein, we address the complexity of the bacterial adaptive response to changing antibiotic selective pressures by studying the long-term *in-patient* evolution of a broad diversity of β-lactam resistant *Pseudomonas aeruginosa* clones. By using mutational and ultra-deep amplicon sequencing, we analyzed multiple generations of a *P. aeruginosa* hypermutator strain persisting for more than 26 years of chronic infection in the airways of a cystic fibrosis (CF) patient. We identified the accumulation of multiple alterations targeting the chromosomally encoded class C β-lactamase (*bla*_PDC_), providing structural and functional protein changes that resulted in a continuous enhancement of its catalytic efficiency and high level of cephalosporin resistance. This evolution was linked to the persistent treatment with ceftazidime, which we demonstrate selected for variants with robust catalytic activity against this expanded-spectrum cephalosporin. Surprisingly, “a gain of function” of collateral resistance towards ceftolozane, a more recently introduced cephalosporin that was not prescribed to this patient, was also observed and the biochemical basis of this cross-resistance phenomenon was elucidated. This work unveils the diversity of evolutionary trajectories driven by bacteria in the natural CF environmental setting, towards a multidrug resistant phenotype after years of antibiotic treatment against a formidable pathogen.

## INTRODUCTION

Our current knowledge regarding the evolution of bacterial antibiotic resistance mainly comes from clinical, microbiological and biochemical studies that are performed under highly controlled conditions (Elena and Lenski, 2003; Weinreich et al., 2006; MacLean et al., 2010; Palmer and Kishony, 2013; Baym et al., 2016; Boolchandani et al., 2019; Card et al., 2021). Collectively, we have learned that the emergence and evolution of antibiotic resistance, one of the greatest challenges to our civilization, is a far more complex phenomenon; few studies exist that offer insights into “real-world” scenarios that adequately or completely explain evolutionary trajectories that shape existing phenotypes (Bershtein et al., 2006; Meini et al., 2015; Stiffler et al., 2015; Prickett et al., 2017; Frimodt-Møller et al., 2018; Andersson et al., 2020; Mehlhoff et al., 2020).

Chronic infections by *Pseudomonas aeruginosa* are main causes of morbidity and mortality in patients suffering from cystic fibrosis (CF). Treating these long-term airway infections is extremely challenging since *P. aeruginosa* displays an intrinsic resistance to many antibiotics, as well as an unwelcome capacity to develop and evolve resistance to newly introduced antibiotics. Acquired antibiotic resistance in CF associated isolates of *P. aeruginosa* occurs mainly through the accumulation of multiple mutations that alter the expression and/or function of different chromosomal genes (Lister et al., 2009; Cabot et al., 2012). Furthermore, *P. aeruginosa* hypermutator strains are frequently isolated from CF patients, thus raising the pace of increased antibiotic resistance development (Oliver et al., 2000; Ciofu et al., 2005; Macia et al., 2005; Montanari et al., 2007; Mena et al., 2008) and the repertoire of adaptability (Lujan et al., 2007; Moyano et al., 2007; Feliziani et al., 2010; Luján et al., 2011; Marvig et al., 2013; Feliziani et al., 2014). Nevertheless, these persistent infections with complex phenotypes offer unique opportunities to study antibiotic resistance evolution since: (i) they are highly influenced by long-term antibiotic treatments to which patients are exposed during their entire lives; (ii) they are often clonal and single *P. aeruginosa* lineages that persist in the lungs of individual patients for many decades; and (iii) *de novo* evolution of antibiotic resistance in individual patients can be monitored constituting an attractive model system for studying the evolution of bacterial populations that strive to adapt to complex dynamic environments (Folkesson et al., 2012).

These pulmonary infections with *P. aeruginosa* in patients with CF are intensively treated with a varied repertoire of antipseudomonal antibiotics, including aminoglycosides, quinolones, and β-lactams. In response, *P. aeruginosa* displays a wide arsenal of resistance mechanisms such as reduced outer membrane permeability, upregulation of multiple broad-spectrum drug efflux pumps, antimicrobial modifying enzymes, and target site changes (Breidenstein et al., 2011). In the case of β-lactam antibiotics, the main resistance mechanism in *P. aeruginosa* is the mutation-mediated overproduction of the chromosomally encoded class C β-lactamase, PDC (*<u>P</u>seudomonas*-<u>d</u>erived <u>c</u>ephalosporinase). Constitutive overexpression of the *ampC* beta-lactamase gene (thereinafter *bla*_PDC_) results from mutations affecting regulatory genes of the peptidoglycan recycling process linked to bacterial cell wall assembly (Moya et al., 2009; Alvarez-Ortega et al., 2010; Tsutsumi et al., 2013; Fisher and Mobashery, 2014; Calvopiña and Avison, 2018). β-lactam resistance has also been associated with structural modifications of PDC (Rodriguez-Martinez et al., 2009; Cabot et al., 2014; Lahiri et al., 2014; Berrazeg et al., 2015; Lahiri et al., 2015; MacVane et al., 2017; Fraile-Ribot et al., 2018), as evidenced by the >400 PDC variants that have been described so far (Oliver, 2020). This impressive number of allelic variants accounts for a highly polymorphic enzyme with a great capacity of tolerating amino acid substitutions, insertions and deletions (Oliver, 2020). Recent studies have shown that clinical resistance to β-lactams is primarily based on specific changes in conserved motifs of PDC, which lead to conformational rearrangements enhancing the catalytic efficiency of the enzyme (Raimondi et al., 2001; Jacoby, 2009; Lahiri et al., 2015; Barnes et al., 2018; Arca-Suárez et al., 2020).

In a whole-genome sequencing study of hypermutator populations of *P. aeruginosa* during long-term chronic airways infections in a single patient (Feliziani et al., 2014), 36 genes in the β-lactam resistome (from a total of 70) carried mutations (Colque et al., 2020). Specifically, the *bla*_PDC_ gene was targeted by multiple independent mutational events in a process boosted by hypermutability (Colque et al., 2020) leading to a wide diversity of coexisting *bla*_PDC_ alleles and high-levels of β-lactam resistance, expanding the range of *P. aeruginosa* opportunities for persistence (Colque et al., 2020). However, the presence of mutations in several genes and the increased expression of PDC compared to isolates preceding the antibiotic treatment does not permit a direct assessment of the impact of the allelic variability of PDC in resistance. Therefore, dissection of the impact of specific mutations in this gene is of relevance to trace the evolution of this enzyme and consequently to guide future therapies.

Here, we unravel the mutational pathways and biochemical mechanisms involved in different PDC variants by analyzing the evolutionary history of more than 25 years of CF chronic infection in a single patient. We show how the combination of substitutions in important amino acid residues in PDC changes the architecture of the enzyme active site. This adaptive scenario led to the selection of distinct enzymatic variants, which conferred resistance to a broad range of β-lactam antibiotics, even to novel combinations of these drugs that were not prescribed to this patient, such as ceftolozane/tazobactam. We also identify the molecular features that elicited the selection of collateral resistance driven by the use of narrow spectrum cephalosporins, and how this resistance was potentiated by hypermutability in *P. aeruginosa*. By modelling a core of three substitutions preserved in the prevailing variants, we hypothesize that favorable interactions are created between ceftolozane and PDC. This work details the trajectory undertaken on the path towards a multidrug resistant phenotype even against untested drugs and illustrates the “collateral damage” suffered by years of antibiotic treatment in the attempt to eradicate this versatile pathogen.

## RESULTS

### Long-term evolution of *P. aeruginosa* hypermutator populations leads to the selection of novel *bla_PDC_* allelic variants with enhanced cephalosporin resistance

In a previous study, the complete genomes of 14 isolates from a hypermutator *P. aeruginosa* lineage spanning 20 years of patient infection history (CFD patient) were sequenced (Feliziani et al., 2014). The clonal collection included a non-mutator ancestor from 1991, two hypermutator isolates from 1995 and 2002, and 11 isolates taken from the same sputum sample in 2011, all of them harboring the same *mutS* mutation (Feliziani et al., 2014). Within-patient genome comparisons revealed a vast accumulation of mutations that shape an extensively diversified population composed of different sub-lineages, which coexisted from the beginning of the chronic infection. Interestingly, the gene encoding the β-lactamase PDC (*bla*_PDC_) was among the most frequently altered by mutations across different isolates, suggesting that *bla*_PDC_ was subjected to strong selective pressure (Colque et al., 2020) (Figure 1). In fact, during the course of chronic airway infection, the patient received prolonged antibiotic treatment with β-lactam antibiotics (Figure 1A). The patient initially received short courses of variable duration of cefotaxime, ceftazidime, piperacillin, aztreonam and the carbapenems, thienamycin and meropenem, and then was continuously treated with ceftazidime from 2004 until the end of 2016. Accordingly, a significant increase in the resistance to ceftazidime in these isolates was reported after the prolonged treatment with this antibiotic (Figure 1-figure supplement 1), which is evident since 2002 (Colque et al., 2020).

**Figure 1.**
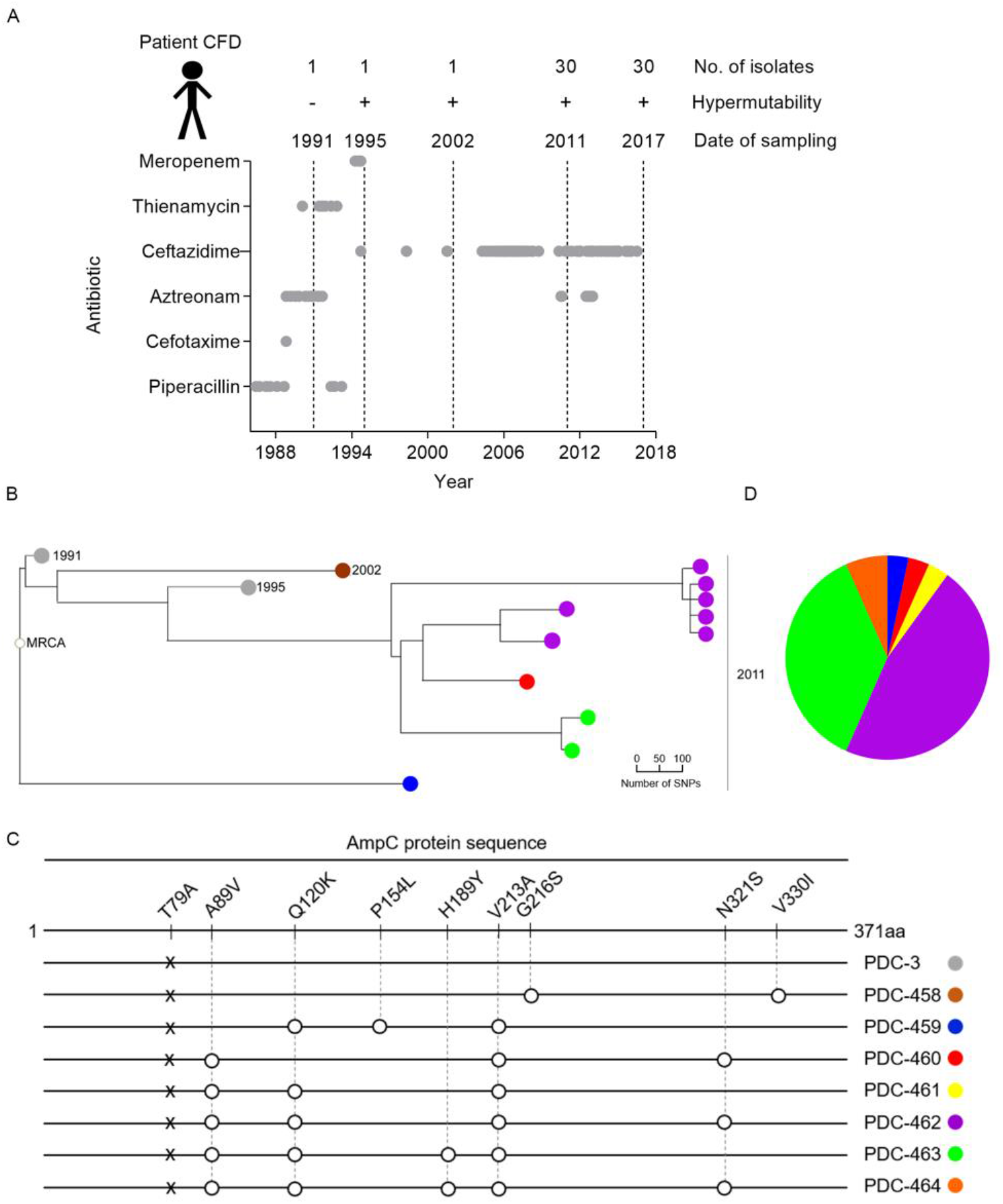
β-lactamase PDC variants from CFD isolates. A) Overview of isolate sampling time points and β-lactam antibiotic treatment throughout the 26-years study. *P. aeruginosa* isolates were collected from patient CFD between 1991 and 2017. Dotted lines indicate the time of isolation of the 1991, 1995, and 2002 isolates, as well as the collections of 30 isolates from single sputum samples in 2011 and 2017. The plus and minus symbols indicate the hypermutability state of the *P. aeruginosa* strains. The β-lactam antibiotics used in chemotherapy are listed on the y axis. Treatment with this group of antibiotics started from 1986 and lasted until end of 2016. Gray circles indicate the start and end of an antibiotic dose. B) Clustering of CFD isolates sequenced in Feliziani *et al*. (Feliziani et al., 2014), based on maximum-parsimony analysis. Circle colors represent different types of PDC variants from CFD isolates. C) Schematic representation of the β-lactamase PDC protein sequence strain PAO1, and of the 8 PDC-variants that emerged during the 20 years of evolution, with their amino acid variations respect to PAO1. Numbering of amino acids refers to the mature protein of PAO1 strain, after cleavage of the 26 N-terminal amino acid residues from the signal peptide, according to the PDC-wide structural position system (SANAC numbering) (Mack et al., 2020). Early isolates from 1991 and 1995 harbor PDC-3 with a T79A polymorphism, which was also present in all the sequenced isolates. Isolate from 2002 harbored PDC-458 variant. The set of 30 isolates evaluated in 2011 harbored PDC-459 to PDC-464 variants. D) Pie chart indicates the percentage of each PDC variant in the 2011 population, respect to the total number of isolates (30) in which *bla*_PDC_ gene was sequenced. PDC-3 differs from PDC-1 from PAO1 by the T79A mutation, which does not affect resistance nor the substrate specificity of the lactamase (Rodríguez-Martínez et al., 2009; Berrazeg et al., 2015).

In an effort to determine the evolution of substrate specificity of the PDC resistant determinant, we tested the resistance of these isolates to ceftolozane-tazobactam, a combination of a cephalosporin and a β-lactamase inhibitor approved in 2014. Even though this combination was not prescribed to this patient, the isolates dating back to 2002 were highly resistant to this combination (Figure 1A-figure supplement 1).

In order to understand the reason for this phenotype, we sequenced *bla_PDC_* from 19 additional isolates belonging to the selected lineage, resulting in a total of 30 isolates obtained from the same 2011 sputum sample. As shown in Figure 1B, the ancestor from 1991 and the 1995 isolate harbored the PDC-3 variant (Rodríguez-Martínez et al., 2009; Berrazeg et al., 2015). After two decades of chronic infection, 7 new *bla_PDC_* allelic variants (referred to as PDC-458 to PDC-466) were identified. Each allele harbored 2 to 5 mutations relative to the ancestral *bla*_PDC-3_, being the result of different combinations of 8 substitutions (Figure 1C).

Substitutions G216S and V330I, which generated variant PDC-458 from the 2002 isolate were not present in the 2011 population, which instead displayed combinations of the other six substitutions distributed in six new PDC variants (Fig 1C). PDC-462 (A89V, Q120K, V213A, N321S) was the most prevalent variant in the 2011 population, being found in 14 (47%) coexisting isolates, followed by variant PDC-463 (A89V, Q120K, H189Y, V213A) that was present in 37% of the isolates. The four remaining allelic variants were rare and only present in one or two isolates (Figure 1D).

### Population dynamics of *bla*_PDC_ mutations during chronic infection

To explore the dynamics of the *bla*_PDC_ allelic prevalence in the population, a new sputum sample from patient CFD was collected in 2017, and a new set of 30 isolates was obtained. Sequencing of the *bla*_PDC_ genes from these isolates revealed a novel scenario (Figure 2A):

**Figure 2.**
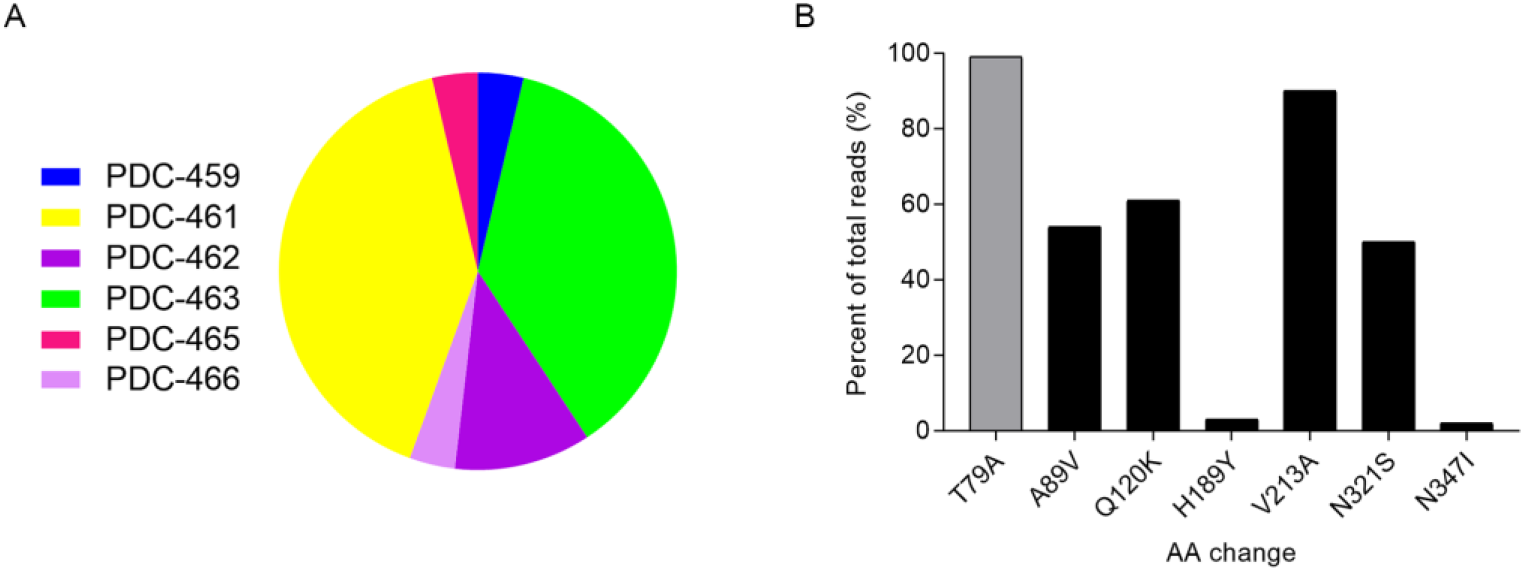
Population analysis on CFD 2017 sample. Sputum sample collected from patient CFD was divided in two and each half was processed for A) Sanger sequencing of *bla*_PDC_ gene in 30 isolates for which sputa was plated in *Pseudomonas* isolation agar and grown for 48hs at 37°C. Pie chart indicates the percentage of each PDC variant in the 2017 population, respect to the total number of isolates sequenced. B) Sequencing of *bla*_PDC_ gene directly amplified on sputum sample for which DNA was extracted and sequenced on Illumina MiSeq. Graph represent the percentage of total reads of each amino acid (AA) variation in the whole population of *P. aeruginosa* (Table supplement 2). T79A (grey bar), considered as a polymorphism that does not affect β-lactam resistance (Berrazeg et al., 2015), was in the 100% of the population confirming CFD patient was colonized by a *P. aeruginosa* lineage derived by a single ancestral clone containing the PDC-3 variant.

PDC-461 (scarcely represented in the 2011 sampling) became prevalent (being present in 39% of the isolates), overriding PDC-463 (36%). Instead, PDC-462, from being a prevalent variant was found in only 11% of the isolates. Two new *bla*_PDC_ alleles, referred to as PDC-465 and PDC-466, were observed. PDC-465 showed the same G216S substitution present in PDC-458, but combined with the novel T256P substitution. On the other hand, PDC-466 seems to derive from PDC-462 through the addition of G205D, thus accumulating five substitutions. These two latter variants displayed a low frequency, like PDC-459, which in 2017 still maintained the low frequency observed in the 2011 population (Figure 2A).

Overall, the main composition of *bla*_PDC_ mutations observed in 2011 was still conserved in the 2017 isolates. We wondered whether these mutations were representative of the whole population. Thus, the 2017 diversity and prevalence of *bla*_PDC_ mutations were analyzed at the population level by performing *bla*_PDC_ amplicon sequencing directly from DNA purified from whole sputum samples obtained from patient CFD (Figure 2B).

Following coverage analysis of >5000 sequencing reads per base in the *bla*_PDC_ open reading frame, only mutations with population frequencies above 2% were considered (Table supplement 2). A89V, Q120K, V213A, N321S were the most frequently observed substitutions, followed by H189Y and N347I (Figure 2B). Interestingly, N347I, which was not observed in any of the isolates previously analyzed, has been reported to confer resistance to cephalosporins (Berrazeg et al., 2015). Substitutions G216S, T256P and G205D, present in PDC-465 and PDC-466, were not detected by this sequencing analysis, probably due to the low prevalence of these variants.

This genetic analysis clearly reveals that substitutions present in the most prevalent PDC variants in either 2011 or 2017 were the most frequent substitutions observed in the global population.

### Combination of multiple *bla*_PDC_ mutations generate resistance to aztreonam and cephalosporins, including ceftolozane

To dissect the role of the mutations present in the different *bla*_PDC_ allelic variants in β-lactam resistance, we designed a system to analyze resistance profiles in a common *Pseudomonas aeruginosa* PAO1 genetic background that allows the control of PDC expression levels. Firstly, a *bla*_PDC_ deficient PAO1 derivative strain (PAΔA) was constructed, in which the chromosomal *bla*_PDC_ gene was deleted. Then, the seven *bla*_PDC_ allelic variants (Figure 1C) together with the ancestor PDC-3 variant from 1991 and the PDC-1 variant from PAO1 strain were cloned into the pMBLe vector under the control of the *lac* operator (González et al., 2016) and transformed into PAΔA. The different variants expressed to similar levels, as revealed by immunodetection (Figure 1C-figure supplement 2, Table supplement 3). Clones of PAΔA carrying different *bla*_PDC_ alleles were challenged against a panel of anti-pseudomonal β-lactam antibiotics commonly used as therapy in CF patients (Table 1).

**Table 1.**
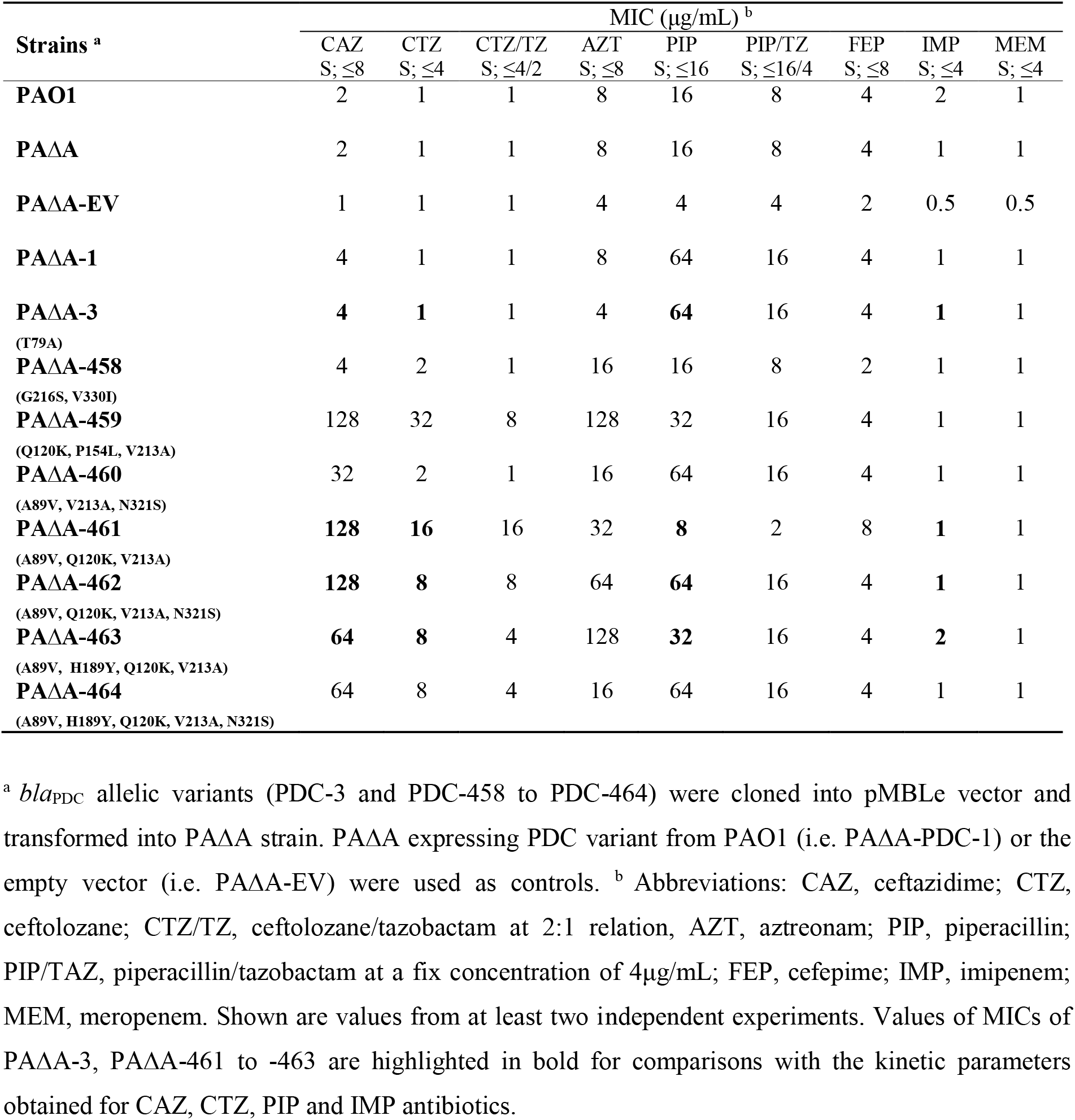
Susceptibility profiles of PAΔA strain complemented with the different PDC variants

Bacteria expressing either PDC-3 or PDC-1 were uniformly susceptible to β-lactams, with the sole exception of piperacillin. In addition, the PDC-458 variant (from the 2002 isolate) resulted in higher resistance levels to aztreonam. In contrast, the most representative allelic variants found in the 2011 isolates (from PDC-459 to PDC-464) showed increased MICs of ceftazidime and aztreonam. These variants present different combinations of A89V, Q120K, H189Y, P154L, V213A, and N321S substitutions, resulting in triple, quadruple and quintuple substitutions. Some of these variants (PDC-459, PDC-461 and PDC-463) showed lower MICs of piperacillin and piperacillin/tazobactam, whereas none of them conferred resistance to cefepime or the carbapenems imipenem and meropenem (Table 1). All variants harboring the Q120K substitution (PDC-459, −461 to −464) showed high resistance levels to ceftolozane (S ≤4 μg/mL), either alone or combined with the β-lactamase inhibitor tazobactam (S ≤4/2 μg/mL) (Table 1). These results show that ceftolozane resistance is due (at least partially) to these substitutions in PDC. The highest resistance to both ceftazidime and ceftolozane was conferred by the PDC-459 combination, Q120K, P154L and V213A followed by PDC-461, which clusters A89V, Q120K and V213A. Additions of N321S and H189Y to the latter triple-mutation combination in PDC-462 and PDC-463, respectively, not only increased resistance to aztreonam but also reverted the decrease in piperacillin resistance observed in PDC-461 (Table 1).

### Differential competitiveness of coexisting-PDC variants can shape the dynamics of resistant subpopulation of *P. aeruginosa* upon exposure to β-lactams

The effect of multiple combined *bla*_PDC_ mutations on bacterial fitness was explored by performing competitive growth assays using the *P. aeruginosa* PAΔA strain carrying *bla*_PDC_ allelic most prevalent variants among CFD 2011 and 2017 populations (referred to as PAΔA-461, PAΔA-462, PAΔA-463, and PAΔA-464), tagged with the *lacZ* gene. These allelic variants combined A89V, Q120K, H189Y, V213A and N321S substitution to generate triple, quadruple or quintuple variant PDCs (Figures 1C and 2). Pairs of tagged/untagged strains were co-cultured *in vitro* and then plated on LB-agar plates containing X-gal.

We first evaluated the relative fitness by competing each variant with the PAΔA strain expressing the ancestral PDC-3 variant (PAΔA-3). As shown in Figure 3A and B, significant differences were not observed in the absence of antibiotics. Instead, in the presence of the β-lactams ceftazidime or aztreonam (the antibiotics used in the therapy of patient CFD), PAΔA-461, PAΔA-462, PAΔA-463, and PAΔA-464 clearly outcompeted PAΔA-0. PAΔA-464 showed lower levels of competitiveness than PAΔA-461, PAΔA-462 and PAΔA-463, indicating that the introduction of a fifth substitution can compromise resistance to these antibiotics.

**Figure 3.**
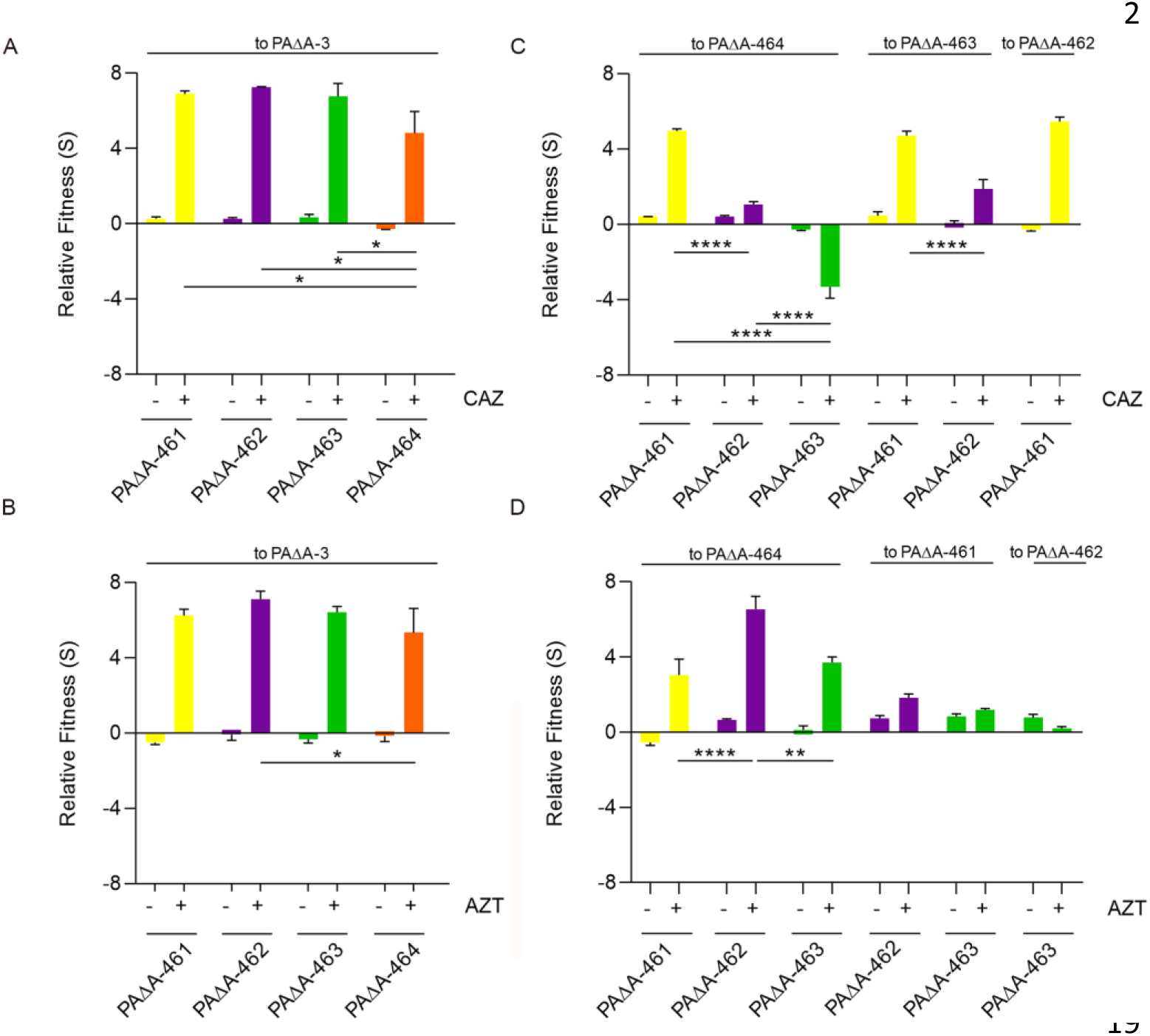
*In vitro* competition experiments among different *bla*_PDC_ alleles. Competition experiments were performed in the absence or presence of ceftazidime (CAZ, A and C) or aztreonam (AZT, B and D). PAΔA strain expressing PDC-461, PDC-462, PDC-463 or PDC-464 (i.e. PAΔA-461, PAΔA-462, PAΔA-463, and PAΔA-464) were competed against PAΔA expressing PDC-3 (PAΔA-3). Fitness (S) relative to PAΔA-3 for (A) ceftazidime or (B) aztreonam are shown. Then, PAΔA-461, PAΔA-462, PAΔA-463 and PAΔA-464 were competed against each other in the absence of antibiotics or presence of (C) ceftazidime or (D) aztreonam. (See SI Appendix for scheme of antibiotic concentration). Measurements were carried out in triplicates for at least two independent experiments, and the results are expressed as means with their SEM. Statistically significant differences at p < 0.0001, p < 0.01 and p < 0.05 are identified by ****, ** and *, respectively (two-way ANOVA followed by Tukey’s Multiple Comparisons Test).

When competed between each other upon ceftazidime exposure, PAΔA-461 showed a clear advantage over PAΔA-462, −463 and −464 (Figure 3C), suggesting that the A89V/Q120K/V213A combination reached the highest relative fitness, whereas additional substitutions impaired competitiveness in the presence of this antibiotic. For instance, the quadruple variant PAΔA-462 showed higher fitness than PAΔA-463 (harboring H189Y instead of N321S) and PAΔA-464 (harboring both, H189Y and N321S), which in turn outcompeted PAΔA-463 (Figure 3C).

In the presence of aztreonam, a clear fitness advantage was observed for all variants against PAΔA-464. Yet, PAΔA-463 and PAΔA-462 showed higher relative fitness than PAΔA-461, suggesting that the addition of N321S and H189Y extend the spectrum of β-lactam resistance (Figure 3D).

These results support that the high prevalence of PDC-462 and PDC-463 variants in the 2011 population are correlated with the simultaneous administration of ceftazidime and aztreonam that took place prior to sample collection (Figs. 1A and D). Subsequently, the repeated rounds of ceftazidime treatments have clearly shaped the 2017 population, where PDC-461, the most resistant variant against cephalosporins, prevailed (Figure 2A).

### PDC variants show improved hydrolytic activity toward ceftazidime and ceftolozane

We next assessed the capacity of the most relevant PDC variants to hydrolyze β-lactams. The mature PDC-3, PDC-461, PDC-462 and PDC-463 proteins were expressed and purified from *E. coli* cultures to homogeneity. Then, we performed steady-state kinetic measurements to test the catalytic efficiencies against the β-lactams ceftazidime, piperacillin and imipenem. In agreement with previous reports (Rodríguez-Martínez et al., 2009; Drawz et al., 2011; Barnes et al., 2018), PDC-3 hydrolyzed efficiently piperacillin while showing a poor hydrolytic activity against ceftazidime, ceftolozane and imipenem (Table 2).

**Table 2.** Kinetic parameters of PDC variants with different substrates

| β-lac<sup>a</sup> | PDC-3 | | | PDC-461 | | | PDC-462 | | | PDC-463 | | |
| --- | --- | --- | --- | --- | --- | --- | --- | --- | --- | --- | --- | --- |
|  | K<sub>M</sub> | k<sub>cat</sub> | k<sub>cat</sub>/K<sub>M</sub> | K<sub>M</sub> | k<sub>cat</sub> | k<sub>cat</sub>/K<sub>M</sub> | K<sub>M</sub> | k<sub>cat</sub> | k<sub>cat</sub>/K<sub>M</sub> | K<sub>M</sub> | k<sub>cat</sub> | k<sub>cat</sub>/K<sub>M</sub> |
| **CAZ** | 57.4<br>±<br>39.3 | 0.0105<br>± 0.0008 | 0.2<br>±<br>0.1 | 93<br>±<br>40 | 0.46<br>±<br>0.01 | 5<br>±<br>2. | 23<br>±<br>2 | 0.12<br>±<br>0.01 | 5.3<br>± 0.9 | 55<br>±<br>27 | 0.13<br>±<br>0.01 | 2.4<br>± 1.3 |
| **PIP** | 19 ±<br>7 | 3<br>±<br>2 | 154<br>±<br>160 | 105<br>±<br>2 | 0.145<br>±<br>0.007 | 1.38<br>± 0.09 | 10<br>±<br>3 | 1.44<br>±<br>0.06 | 146<br>± 5 | 54.5<br>±<br>3 | 0.8<br>±<br>0.2 | 15<br>± 1 |
| **IMP** | 10.4<br>±<br>3 | 0.016±<br>0.01 | 2<br>±<br>1 | 3±<br>2 | 0.007<br>±<br>0.002 | 0.2<br>±<br>0.2 | 37<br>±<br>4 | 0.039<br>±<br>0.005 | 1<br>± 0.2 | 36<br>±<br>5 | 0.013<br>±<br>0.001 | 0.36<br>± 0.09 |
| **CTZ<sup>b</sup>** | ND | ND | 0.04<br>± 0.01 | ND | ND | 6.1<br>±<br>1.3 | ND | ND | 0.7<br>± 0.1 | ND | ND | 0.33<br>± 0.06 |

<sup>a</sup>Kinetic parameters were determined for ceftazidime (CAZ), piperacillin (PIP), imipenem (IMP)
and ceftolozane (CTZ). Two independent experiments were carried out and results are expressed
as means. Units: K<sub>m</sub> (μM). k<sub>cat</sub> (s<sup>-1</sup>) and k<sub>cat</sub>/K<sub>m</sub> (mM<sup>-1</sup> s<sup>-1</sup>). <sup>b</sup> Due to the low hydrolysis of
ceftolozane, k<sub>cat</sub>/K<sub>m</sub> values were obtained from Lineweaver-Burk plots. ND, not determined.

PDC-461, −462 and −463 efficiently hydrolyzed ceftazidime, showing 28-, 29- and 13-fold increased catalytic efficiencies (respectively) relative to the ancestor PDC-3, indicating that mutations in these PDC improved their catalytic performance on this cephalosporin. PDC-461 displayed a catalytic efficiency against piperacillin 100-fold impaired with respect to PDC-3, disclosing a tradeoff in the substrate profile shaped by the presence of the three core mutations (A89V, Q120K and V213A). Instead, the additional mutations present in PDC-462 and PDC-463 were able to restore this activity. Indeed, PDC-462 displayed hydrolytic levels against piperacillin similar to PDC-3, in agreement with the observed piperacillin MICs for the strain expressing this variant (Table 1).

All PDC variants maintained low hydrolysis rates for imipenem, displaying *k*_cat_ values between 0.01 and 0.04 s^−1^, which correlate well with the imipenem susceptibility (MICs of 1 to 2 μg/mL) observed in PAΔA expressing either PDC-461, −462 or −463 variants (Table 1). Remarkably, when the catalytic efficiencies of PDC-461, −462 and −463 were assessed against the recently introduced cephalosporin ceftolozane, the *k*_cat_/*K*_m_ ratios showed significantly increased values compared to that of the parental enzyme. PDC-462 and 463 show 16- and 8-fold enhancements of this activity, a performance that is largely overcome by PDC-461, showing a 150-fold increase in *k*_cat_/*K*_m_.

These catalytic efficiencies correlate very well with the MIC levels of different antibiotics determined in an isogenic *Pseudomonas* background (Table 1), revealing that the different accumulated mutations are responsible of tuning the substrate profile of these variants.

### Molecular dynamics simulations reveal enlargement of substrate-binding pocket in PDC evolved mutants

The different substitutions present in the studied variants are scattered in the protein structure, many of them being part of protein loops (Figure 4). Expansion of the substrate profile by mutations in β-lactamases has been accounted for by changes in the protein dynamics. Therefore, we performed classical molecular dynamics (MD) simulations on these variants (PDC-3, PDC-461, PDC-462, and PDC-463) in the unbound state. MD simulations were run for 200 ns and conformational clustering was performed as described (González et al., 2014; Morán-Barrio et al., 2016; González et al., 2018). All proteins preserved their global tertiary structure during the MD simulations but revealed significant changes in the substrate binding pocket elicited by the substitutions (Figure 5).

**Figure 4.**
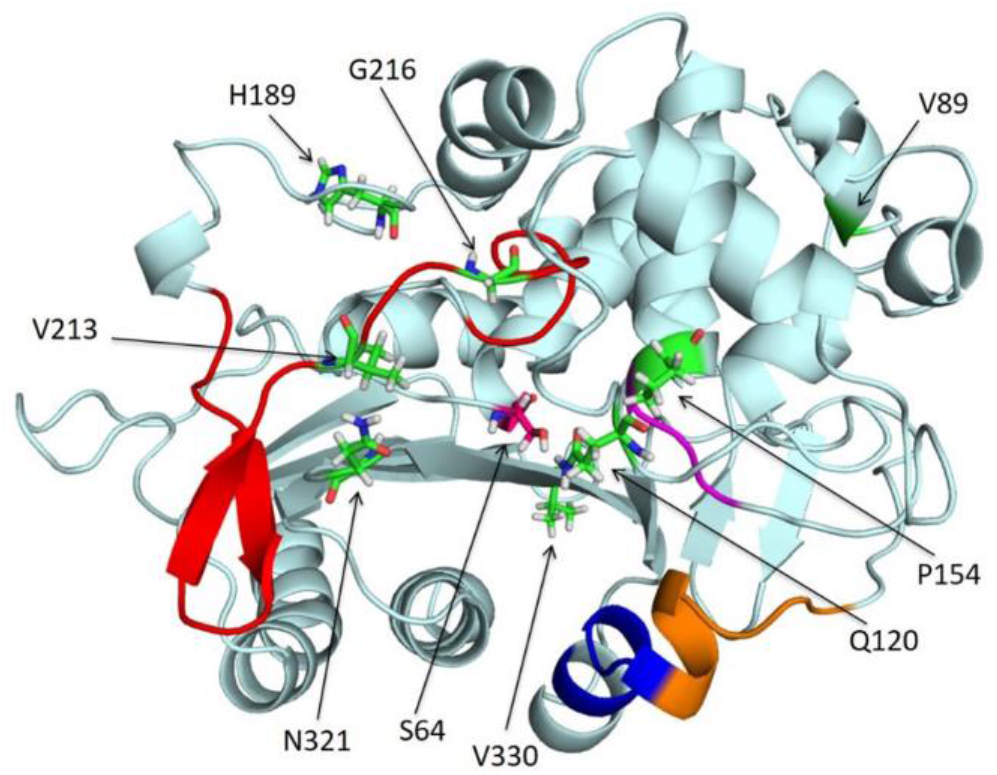
Representation of the PDC β-lactamase structure from *P. aeruginosa* PAO1 (PDB 4OOY doi:10.2210/PDB4OOY/PDB). The different structural regions lining the binding site are colored as follows: omega loop, red; helix H-10, blue; R2 loop, orange; and YSN, purple. The amino acids residues, which were mutated across the different *bla*_PDC_ allelic variants in this study are represented with sticks in green and pointed with arrows.

**Figure 5.**
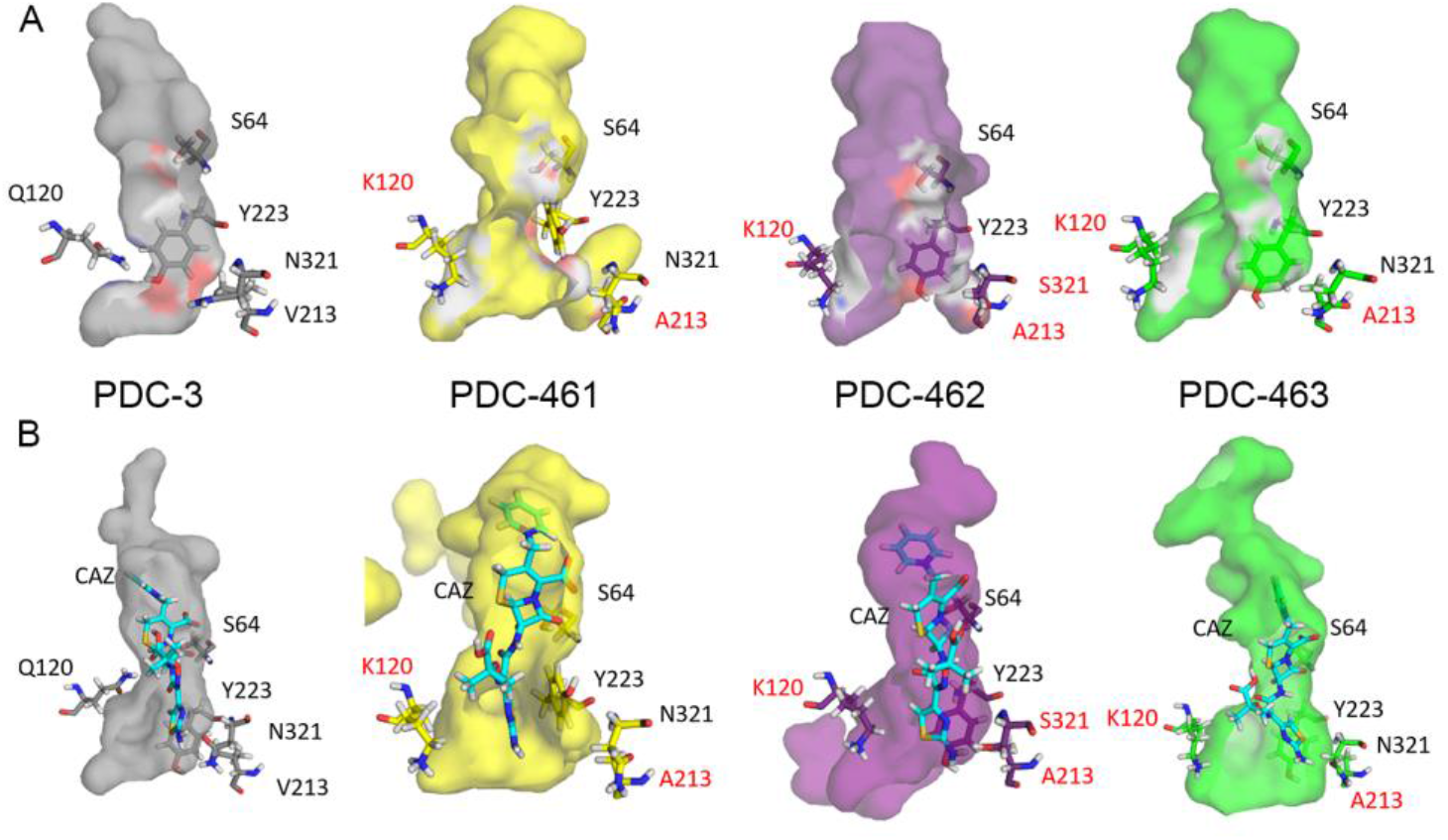
Molecular modeling of PDC proteins. Structures of the apo/free versions (A) and coupled with ceftazidime antibiotic CAZ (B) with their active site cavities are shown. Colors of protein structures: PDC-3 (grey), PDC-461 (yellow), PDC-462 (purple) and PDC-463 (green).

The three variants showing an enhanced activity towards ceftazidime and ceftolozane (PDC-461, PDC-462, and PDC-463) present a broader active site cavity (Figure 5B-figure supplement 3). Substitution V213A induces a conformational change in the Ω-loop, involving residues 200-223. As a result, a hydrogen bond formed by phenolic OH of Y223 with the backbone of G214 present in PDC-3 is lost, inducing a conformational change in Y223 in PDC-461. Mutation Q120K eliminates a hydrogen bonding interaction of the amide side chain with N153 (from the YSN loop, located in the opposite side of the substrate binding pocket). This results in a conformational change of this residue, with Lys120 pointing outwards and therefore further widening the active site cavity (Figure 6). The structural impact of this mutation is similar in PDC-461, −462 and −463, i.e., regardless the genetic background, highlighting the key role of the Q120K mutation in the evolution of resistance.

**Figure 6.**
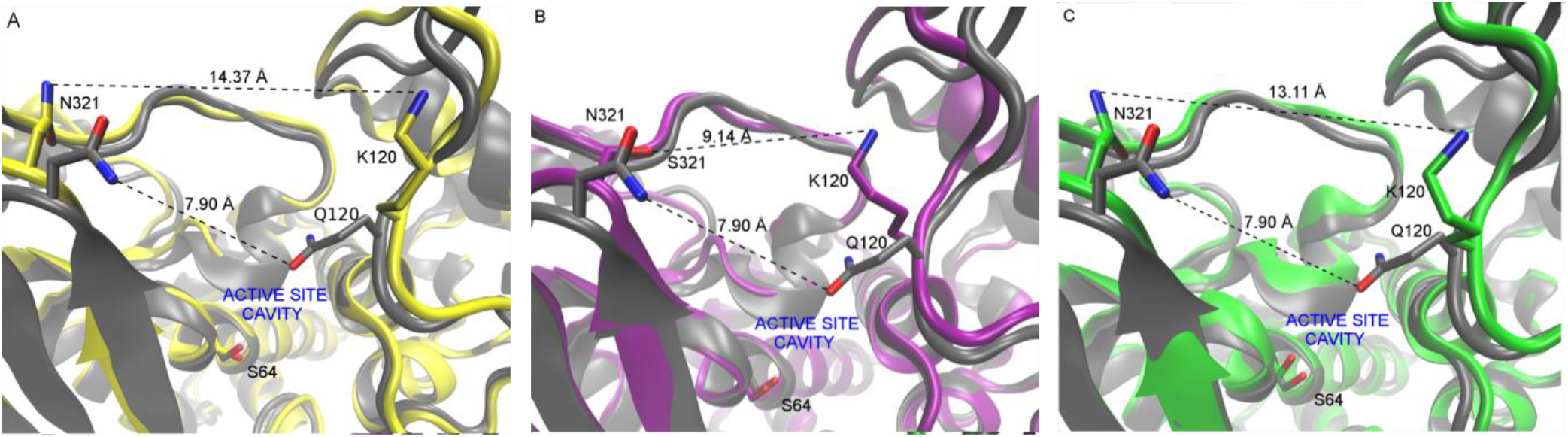
Comparison of representative snapshots of the MD simulations of each protein studied vs PDC-3 showing the active site cavity environment (R1 region). In all representations, N atoms are depicted in blue, oxygen atom in red and key residues are highlighted in sticks. C atoms of the PDC-3 are depicted in grey. (A) PDC-3 vs PDC-461 (C atoms in yellow); (B) PDC-3 vs PDC-462 (C atoms in purple); (C) PDC-3 vs PDC-463 (C atoms in green). Residue distances are depicted with dashed lines.

Representative conformations extracted from the MD simulations were used to build *in silico* the complexes of each PDC variant with ceftazidime, using the crystal structure of acylated ceftazidime bound to EDC (*<u>E</u>scherichia coli*-<u>d</u>erived <u>c</u>ephalosporinase) (PDB 1IEL) (Powers et al., 2001) as a template. In order to understand the interaction of ceftazidime with the protein environment, we optimized these structures by hybrid quantum mechanics-molecular mechanics (QM-MM) simulations. In all four complexes, ceftazidime interacts with residues S64, K67, N153, Y223, S319, N321, N344, and N347, in agreement with the previous structural information (Powers et al., 2001; Barnes et al., 2018). The active site changes in the mutants allows a better accommodation of the bulky R1 side chain from ceftazidime (Figure 5B-figure supplement 3). In addition, the conformational change of Y223 results in an aromatic stacking interaction with the 4-thiazolyl ring of ceftazidime at the R1 substituent (Figure 7). The N321S substitution in PDC-462 removes a hydrogen bond with A213 at the Ω-loop that enables to recover the interaction between Y223 and G214. Overall, all variants show a broadening of the active site that accounts for the large increase in activity and resistance against ceftazidime of these PDC variants.

**Figure 7.**
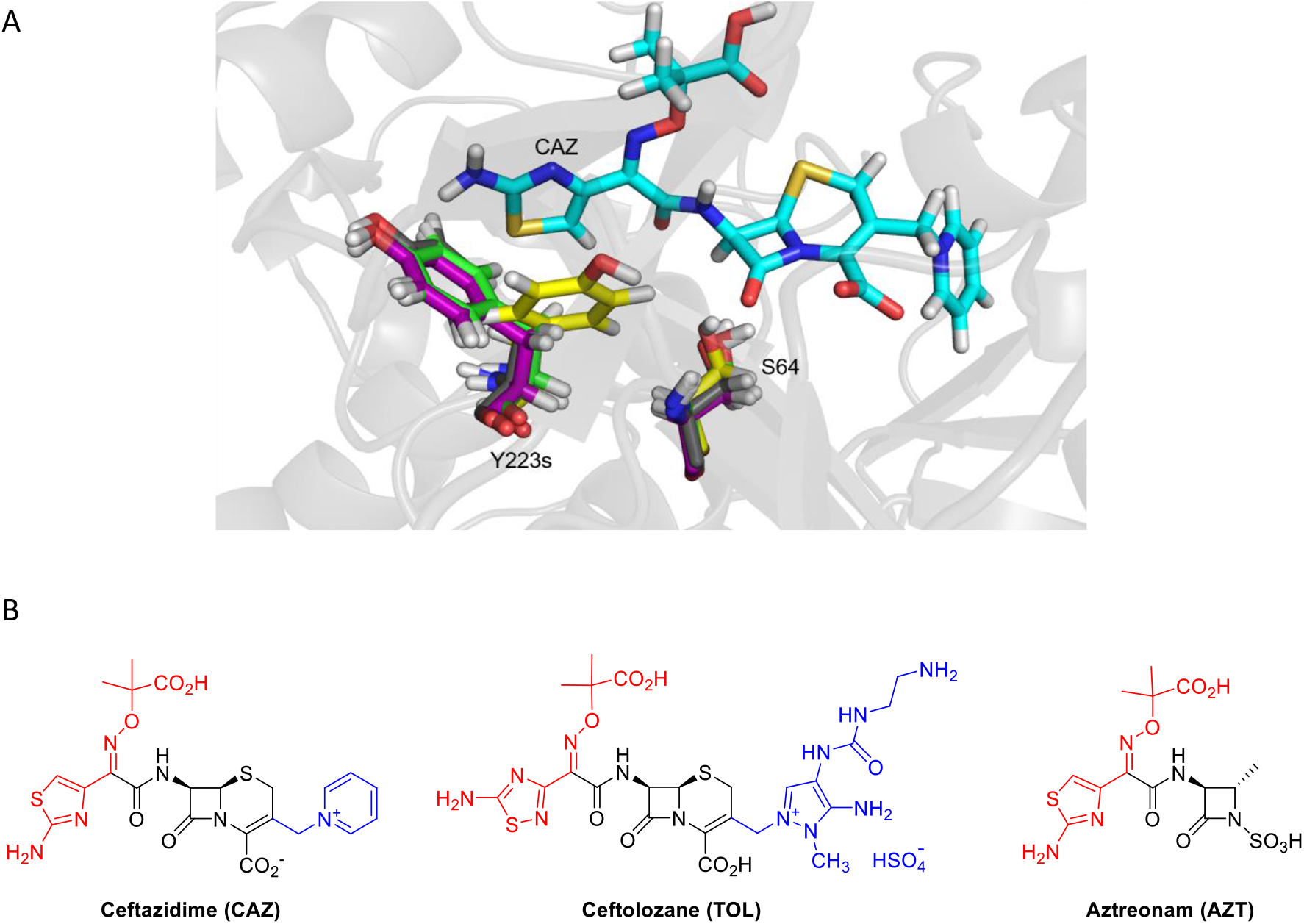
Orientation change in Y223 residue and evidence of a stacking interaction. (A) When ceftazidime (CAZ) antibiotic was introduced in PDC protein structures modeling, a strong orientation change in Y223 residue of PDC-461 was seen. Y223 residues are colored in gray, yellow, purple and green as for their PDC variants PDC-3, PDC-461, PDC-462 and PDC-463 respectively. Structure of CAZ antibiotic and the location of S64 active site is also depicted. (B) Structures of β-lactam antibiotics for which PDC variants showed increased resistance. The R1 side chains of the β-lactam antibiotics are shown in red, and the R2 side chains of the cephalosporins and monobactam aztreonam are shown in blue.

## DISCUSSION

Laboratory-based experimental evolution experiments have been extensively used to replicate bacterial adaptation during antibiotic therapy (MacLean et al., 2010; Palmer and Kishony, 2013; Cabot et al., 2014; Baym et al., 2016; Card et al., 2019; Windels et al., 2020; Card et al., 2021). This approach reduces complexity in order to establish reproducible and well-controlled conditions, from which evolutionary changes and mutational trajectories can be analyzed. However, it has become increasingly clear that *in vitro* antibiotic selection experiments do not necessarily replicate resistance development taking place in more complex settings, where many variables and selective factors can influence the adaptive potential and outcome in bacterial populations (Didelot et al., 2016; Baker et al., 2018). Exploring long-term evolution in ‘natural’ environments, such as those from within-host populations, allows one to embrace this complexity, providing new information that can be relevant for the development of novel strategies to control and/or prevent antibiotic resistance during infections. Here we investigated resistance as a consequence of bacterial evolution in presence of all the patient co-factors (tissues, immune responses, microbiota compositions, etc.), taking the experimental evolutionary system that is exactly what creates the society associated resistance problem: the antibiotic-treated patient.

A previous study of the evolution of *P. aeruginosa* hypermutator lineages combining longitudinal and cross-sectional analysis covering decades of CF chronic infection showed that antibiotic resistance increases as infection progresses towards the establishment of a highly diversified population, that converges towards multidrug resistance (Feliziani et al., 2014; Colque et al., 2020). Here we have dissected the specific effects of mutations accumulated in the β-lactamase *ampC* gene (*bla*_PDC_) in isolates from a single patient who received a long-term treatment (26 years) with β-lactam antibiotics. Importantly, the adaptive evolution of PDC resulted from an accumulation of multiple mutations in the *bla*_PDC_ gene that, when combined, resulted in high level β-lactam resistance. In particular, we show a large increase in the hydrolytic capability of variants PDC-461, PDC-462 and PDC-463 against ceftazidime and ceftolozane, providing a structural and functional rationale, and analyze these findings in the framework of the complexity of the bacterial population elicited by a hypermutator strain.

The concept of population phenotype is related to the frequencies and distribution of relevant alleles in bacterial populations, which can shape community functions and influence clinical outcomes (Azimi et al., 2020). In this work, we demonstrate how the dynamics of multiple and changing antibiotic pressures, together with higher mutation rates, resulted in high allelic variations of the PDC β-lactamase in a diversified resistant clonal population that originally evolved from a susceptible ancestral infecting strain. Noteworthy, the many coexisting PDC variants, each conferring particular spectrum resistance profiles, provide a wider β-lactam resistant phenotype to the population as a whole. For example, one of the most prevalent variants, PDC-461 confers the highest resistance to cephalosporins in an activity trade-off that causes collateral sensitivity to piperacillin. During subsequent steps along therapy, PDC-462 and PDC-463 have acquired insertions of novel “gain-of-function” substitutions in the PDC-461 background, which result in a wider substrate spectrum maintaining cephalosporin resistance, with higher resistance to aztreonam, and restored resistance to piperacillin.

The high efficiency acquired by these PDC variants to confer resistance to the novel anti-pseudomonal cephalosporin, ceftolozane, is of great interest. To our knowledge, this is the first study describing collateral resistance to ceftolozane in a patient that has never been treated with this antibiotic. Resistance to ceftolozane has previously been observed in *P. aeruginosa* infected patients when treated with this antibiotic (Munita et al., 2017; Fraile-Ribot et al., 2018), and other studies have demonstrated that expression of a PDC-3 variant carrying a single E221K mutation can confer high MICs of ceftolozane in *E. coli* (Barnes et al., 2018). It has also been shown that a substitution of Asp219 at the Ω-loop 219, selected after treating a multi-drug resistant *P. aeruginosa* strain with ceftolozane/tazobactam, enhances hydrolysis of this cephalosporin/β-lactamase inhibitor combination *in vivo* (Arca-Suárez et al., 2020). However, *in vitro* long-term experiments of wild-type and mutator strains of *P. aeruginosa* exposed to increasing concentrations of ceftolozane/tazobactam, showed that only mutator strains were able to develop high-levels of resistance, by acquiring multiple mutations that led to overexpression and structural modifications of PDC (Cabot et al., 2014). The value of an increased mutational and innovative potential caused by hypermutability is documented in our *in-patient* study, making it understandable why hypermutators are frequently isolated from bacterial populations challenged by stressful conditions over long time periods (Oliver et al., 2000; Denamur and Matic, 2006; Matic, 2019).

In addition to diverse polymorphisms previously described for PDC, we report three new amino acid substitutions: A89Y, G205D and T256P, which together with Q120K, P154L, H189Y, V213A, G216S, N321S, V330I and N347I mutations generated novel *bla*_PDC_ allelic variants, each harboring from 2 to 5 mutations. P154L, V213A, and N347I are located next to the conserved YSN loop, the C-terminal region of Ω-loop and the C3/C4 recognition region, respectively, and have been shown to individually confer resistance to β-lactam antibiotics in *P. aeruginosa* clinical isolates (Berrazeg et al., 2015; Barnes et al., 2018). The finding of new combinations that further boost cephalosporin resistance supports the adaptability of the PDC scaffold to tolerate various mutations, which at the same time provides a substantial gain-of-function.

The three most prevalent alleles are those that combine mutations A89V, Q120K and V213A and, at the same time, confer the highest resistance to cephalosporins and enhanced competitiveness in the presence of ceftazidime. Molecular dynamics simulations revealed that these three mutations give rise to a wider substrate-binding pocket in the PDC variants. This change favors binding of ceftazidime to the active site, providing space for better accommodating the R1 side chain of this antibiotic (Figure 7A). In addition, substitutions Q120K and V213A trigger a different orientation in the aromatic group of Y223 in the PDC-461 variant, favoring a stacking interaction with the 4-thiazolyl ring present in the R1 group. Other substitutions located in or near the Ω-loop have been shown to enhance cephalosporin resistance by altering the conformation of Y223 (Powers et al., 2001; Thomas et al., 2010; Barnes et al., 2018). This conformational change improves hydrolysis of ceftazidime, ceftolozane and aztreonam, but has the opposite effect on piperacillin. Instead, the N321S substitution in variant PDC-462 restores the Y223 orientation present in PDC-3 while maintaining the enlargement of the substrate-binding pocket, thus extending the hydrolysis towards ceftazidime, ceftolozane, aztreonam and piperacillin. These results suggest that the broadening of the active site induced by the different mutations is more relevant than the interaction with Y223.

The Q120K mutation results in a net widening of the active site entrance, due to the conformational change of the side chain (Figure 6). The impact of Q120K in resistance is evident from analyzing PDC-462 and PDC-460, which differ only by this substitution. The presence of this substitution elicits an impressive increase by 3-fold dilutions in the MICs towards ceftazidime, aztreonam, and ceftolozane (Table 1). We conclude that Q120K plays an important role in the evolution of resistance in this family of variants.

These structural changes result in a better accommodation of the R1 group from ceftazidime: the volume in the active site region that recognizes R1 is increased by more than two-fold in PDC-461, PDC-462 and PDC-463 compared to PDC-3. Interestingly, the R1 group structure, which has been associated with the antipseudomonal activity (Zasowski et al., 2015), is almost identical in aztreonam, ceftazidime, and ceftolozane (Figure 7B). In contrast, changes in the volume of the active site cleft where R2 is located are not perceived to be important for the binding of ceftazidime. We conclude that optimization of R1 binding driven by ceftazidime circumvents this problem and provides a better recognition of both aztreonam and ceftolozane. In the report by Berrazeg and coworkers (Berrazeg et al., 2015), the authors proposed that either ceftazidime or cefepime could induce this effect. In light of the current study, we conclude that it is unlikely that cefepime could elicit a similar cross-resistance effect as ceftazidime since the R1 group is different in this antibiotic.

In conclusion, this study combines genetic, biochemical and structural analyses of the evolutionary process of antibiotic resistance in the natural environment where it truly occurs. This detailed scrutiny of the evolution of a *P. aeruginosa* clone persisting within a single patient reveals how the consistent and intensive antibiotic treatment in the setting of a hypermutator genotype leads to a multidrug resistance phenotype, primarily driven by combined substitutions in the chromosomal β-lactamase, PDC. The amazing plasticity of the PDC structure not only confirms the already known capacity to evolve when facing the challenge of new β-lactams, but also warns us that chemical similarities among β-lactams from different generations could lead to an unexpected evolution of resistance, particularly in the context of a chronic infection by a hypermutator strain. Altogether, our results emphasize the huge evolutionary potential of hypermutator strains and uncover the link between the antibiotic prescription history and the *in-patient* evolution of antibiotic resistance that relies on a molecular-based hypothesis of the adaptation of the PDC β-lactamase. Finally, they highlight the importance of integrating bench-to-bedside research to fully understand the processes that lead to antibiotic resistance.

## MATERIALS AND METHODS

Clinical isolates were obtained from sputum samples from an adult patient with cystic fibrosis attending the Copenhagen Cystic Fibrosis Center at University Hospital Rigshospitalet, Denmark (CFD Patient) (Feliziani et al., 2014). The use of the samples was approved by the local ethics committee of the Capital Region of Denmark (Region Hovedstaden; registration numbers H-A-141 and H-1-2013-032), and patient gave informed consent.

Isolation and identification of *P. aeruginosa* from sputum was carried out as previously described (Høiby and Frederiksen, 2000). Patient age at the time of the first isolate collection was 23 years and the onset of the chronic infection with *P. aeruginosa* was in 1986. The *P. aeruginosa* collection included: an initial isolate from 1991, two intermediate isolates from 1995 and 2002, and two populations of isolates collected in 2011 (Feliziani et al., 2014) and 2017 (this study). For cross-sectional analysis, 30 isolates were taken randomly from the 2011 and 2017 sputum samples. Isolates were stored at −70°C in glycerol stock solution. The 2017 sputum sample was divided in two: one for the isolation of *P. aeruginosa* clones and the other for DNA extraction for ultra-deep sequencing analysis.

The sequences of the *bla*_PDC_ gene corresponding to the described PDCs variants have been deposited in GenBank under the accession numbers shown in SI Appendix.

Materials and methods describing the sequence analysis of *bla*_PDC_ gene in *P. aeruginosa* CFD isolates, DNA extraction and PCR amplification of *bla*_PDC_ gene from whole sputum samples; the construction of *P. aeruginosa* Δ*bla*_PDC_ deficient (PAΔA) and *bla*_PDC_-*lacZ* (PAΔA-*lacZ*) strains; cloning of *bla*_PDC_ allelic variants and expression levels of PDCs in pMBLe vector; competition experiments for determination of competitive fitness of *bla*_PDC_ variants; expression and purification of PDC proteins; classical molecular dynamic simulations and QM-MM calculations with the PDC variants (in apo versions) and PDC in complex with ceftazidime are described in detail in the SI Appendix section Materials and Methods.

## ACKNOWLEDGEMENTS

This work was supported by ANPCyT (Grant N° PICT-2016-1545 and PICT-2019-1590 to AMS, PICT-2016-1657 to AJV, PICT-2019-1358 to PET and PICT-2016-1926 to AAO); SECYT-UNC (Grant N° 33620180100413CB to AMS); MINCyT-Córdoba (Grant N° PID-2018-Res 144 to AMS); NIH (Grant N° R01AI100560 to AJV); the Novo Nordisk Foundation (Grant N° NNF12OC1015920, NNF15OC0017444 and NNF18OC0052776 to HKJ); Rigshospitalet Rammebevilling 2015-17 (Grant N° R88-A3537 to HKJ); Lundbeckfonden (Grant N° R167-2013-15229 to HKJ); Det Frie Forskningsråd FSS (Grant N° DFF-4183-00051 to HKJ); RegionH rammebevilling and Savværksejer Jeppe Juhl og Hustru Ovita Juhls Memorial Fund (Grant N° R144-A5287 to HKJ); and the National Institute of Allergy and Infectious Diseases of NIH (Grant N° R01AI100560, R01AI063517, and R01AI072219 to RAB). A grant provided by Merck & Co., Inc., Kenilworth, NJ USA and the Cleveland Department of Veterans Affairs supported RAB (Grant N° 1I01BX001974) from the Biomedical Laboratory Research & Development Service of the VA Office of Research and Development, and the Geriatric Research Education and Clinical Center VISN 10. The content is solely the responsibility of the authors and does not necessarily represent the official views of the NIH or the Department of Veterans Affairs. AMS, AJV, PET, AAO and AJM are staff members from CONICET. CAC, GD and LGH are recipients of fellowships from CONICET, Argentina.

## AUTHOR CONTRIBUTIONS

AMS and AJV designed research and supervised the study. CAC, PET, AGAO, GD, RAH, LGH, SF, AJM performed experimental research. HKJ provided clinical samples and bacterial collection. CAC and LMS analyzed bioinformatics ultra-deep sequencing data. DMM, performed molecular modeling analyses. CAC, PET, AGAO, DMM, RAB, HKJ, SM, AJV and AMS analyzed data; RAB, SM, AJV and ASM wrote the paper.

## REFERENCES

Alvarez-Ortega, C., Wiegand, I., Olivares, J., Hancock, R.E., and Martínez, J.L. (2010) Genetic determinants involved in the susceptibility of *Pseudomonas aeruginosa* to β-lactam antibiotics. Antimicrobial agents and chemotherapy 54: 4159–4167.

Andersson, D.I., Balaban, N.Q., Baquero, F., Courvalin, P., Glaser, P., Gophna, U. et al. (2020) Antibiotic resistance: turning evolutionary principles into clinical reality. FEMS Microbiology Reviews 44: 171–188.

Arca-Suárez, J., Vázquez-Ucha, J.C., Fraile-Ribot, P.A., Lence, E., Cabot, G., Martínez-Guitián, M. et al. (2020) Molecular and biochemical insights into the in vivo evolution of AmpC-mediated resistance to ceftolozane/tazobactam during treatment of an MDR *Pseudomonas aeruginosa* infection. J Antimicrob Chemother.

Azimi, S., Roberts, A.E.L., Peng, S., Weitz, J.S., McNally, A., Brown, S.P., and Diggle, S.P. (2020) Allelic polymorphism shapes community function in evolving *Pseudomonas aeruginosa* populations. The ISME Journal 14: 1929–1942.

Baker, S., Thomson, N., Weill, F.-X., and Holt, K.E. (2018) Genomic insights into the emergence and spread of antimicrobial-resistant bacterial pathogens. Science 360: 733–738.

Barnes, M.D., Taracila, M.A., Rutter, J.D., Bethel, C.R., Galdadas, I., Hujer, A.M. et al. (2018) Deciphering the evolution of cephalosporin resistance to ceftolozane-tazobactam in *Pseudomonas aeruginosa*. mBio 9: e02085–02018.

Baym, M., Stone, L.K., and Kishony, R. (2016) Multidrug evolutionary strategies to reverse antibiotic resistance. Science 351: aad3292.

Berrazeg, M., Jeannot, K., Ntsogo Enguene, V.Y., Broutin, I., Loeffert, S., Fournier, D., and Plesiat, P. (2015) Mutations in beta-lactamase AmpC increase resistance of *Pseudomonas aeruginosa* isolates to antipseudomonal cephalosporins. Antimicrob Agents Chemother 59: 6248–6255.

Bershtein, S., Segal, M., Bekerman, R., Tokuriki, N., and Tawfik, D.S. (2006) Robustness–epistasis link shapes the fitness landscape of a randomly drifting protein. Nature 444: 929–932.

Boolchandani, M., D’Souza, A.W., and Dantas, G. (2019) Sequencing-based methods and resources to study antimicrobial resistance. Nature Reviews Genetics 20: 356–370.

Breidenstein, E.B., de la Fuente-Nunez, C., and Hancock, R.E. (2011) *Pseudomonas aeruginosa*: all roads lead to resistance. Trends Microbiol 19: 419–426.

Cabot, G., Bruchmann, S., Mulet, X., Zamorano, L., Moya, B., Juan, C. et al. (2014) *Pseudomonas aeruginosa* ceftolozane-tazobactam resistance development requires multiple mutations leading to overexpression and structural modification of AmpC. Antimicrob Agents Chemother 58: 3091–3099.

Cabot, G., Ocampo-Sosa, A.A., Dominguez, M.A., Gago, J.F., Juan, C., Tubau, F. et al. (2012) Genetic markers of widespread extensively drug-resistant *Pseudomonas aeruginosa* high-risk clones. Antimicrob Agents Chemother 56: 6349–6357.

Calvopiña, K., and Avison, M.B. (2018) Disruption of *mpl* Activates β-Lactamase Production in *Stenotrophomonas maltophilia* and *Pseudomonas aeruginosa* Clinical Isolates. Antimicrobial Agents and Chemotherapy 62: e00638–00618.

Card, K.J., LaBar, T., Gomez, J.B., and Lenski, R.E. (2019) Historical contingency in the evolution of antibiotic resistance after decades of relaxed selection. PLOS Biology 17: e3000397.

Card, K.J., Thomas, M.D., Graves, J.L., Barrick, J.E., and Lenski, R.E. (2021) Genomic evolution of antibiotic resistance is contingent on genetic background following a long-term experiment with *Escherichia coli*. Proceedings of the National Academy of Sciences 118: e2016886118.

Ciofu, O., Riis, B., Pressler, T., Poulsen, H.E., and Hoiby, N. (2005) Occurrence of hypermutable *Pseudomonas aeruginosa* in cystic fibrosis patients is associated with the oxidative stress caused by chronic lung inflammation. Antimicrob Agents Chemother 49: 2276–2282.

Colque, C.A., Albarracín Orio, A.G., Feliziani, S., Marvig, R.L., Tobares, A.R., Johansen, H.K. et al. (2020) Hypermutator *Pseudomonas aeruginosa* exploits multiple genetic pathways to develop multidrug resistance during long-term infections in the airways of cystic fibrosis patients. Antimicrobial Agents and Chemotherapy: AAC.02142-02119.

Denamur, E., and Matic, I. (2006) Evolution of mutation rates in bacteria. Molecular Microbiology 60: 820–827.

Didelot, X., Walker, A.S., Peto, T.E., Crook, D.W., and Wilson, D.J. (2016) Within-host evolution of bacterial pathogens. Nature Reviews Microbiology 14: 150–162.

Drawz, S.M., Taracila, M., Caselli, E., Prati, F., and Bonomo, R.A. (2011) Exploring sequence requirements for C3/C4 carboxylate recognition in the *Pseudomonas aeruginosa* cephalosporinase: Insights into plasticity of the AmpC β-lactamase. Protein science : a publication of the Protein Society 20: 941–958.

Elena, S.F., and Lenski, R.E. (2003) Evolution experiments with microorganisms: the dynamics and genetic bases of adaptation. Nature Reviews Genetics 4: 457–469.

Feliziani, S., Marvig, R.L., Luján, A.M., Moyano, A.J., Di Rienzo, J.A., Krogh Johansen, H. et al. (2014) Coexistence and within-host evolution of diversified lineages of hypermutable *Pseudomonas aeruginosa* in long-term cystic fibrosis infections. PLOS Genetics 10: e1004651.

Feliziani, S., Luján, A.M., Moyano, A.J., Sola, C., Bocco, J.L., Montanaro, P. et al. (2010) Mucoidy, quorum sensing, mismatch repair and antibiotic resistance in *Pseudomonas aeruginosa* from cystic fibrosis chronic airways infections. PLOS ONE 5: e12669.

Fisher, J.F., and Mobashery, S. (2014) The sentinel role of peptidoglycan recycling in the beta-lactam resistance of the gram-negative *Enterobacteriaceae* and *Pseudomonas aeruginosa*. Bioorg Chem 56: 41–48.

Folkesson, A., Jelsbak, L., Yang, L., Johansen, H.K., Ciofu, O., Høiby, N., and Molin, S. (2012) Adaptation of *Pseudomonas aeruginosa* to the cystic fibrosis airway: an evolutionary perspective. Nature Reviews Microbiology 10: 841–851.

Fraile-Ribot, P.A., Cabot, G., Mulet, X., Perianez, L., Martin-Pena, M.L., Juan, C. et al. (2018) Mechanisms leading to *in vivo* ceftolozane/tazobactam resistance development during the treatment of infections caused by MDR *Pseudomonas aeruginosa*. J Antimicrob Chemother 73: 658–663.

Frimodt-Møller, J., Rossi, E., Haagensen, J.A.J., Falcone, M., Molin, S., and Johansen, H.K. (2018) Mutations causing low level antibiotic resistance ensure bacterial survival in antibiotic-treated hosts. Scientific Reports 8: 12512.

González, L.J., Moreno, D.M., Bonomo, R.A., and Vila, A.J. (2014) Host-Specific Enzyme-Substrate Interactions in SPM-1 Metallo-β-Lactamase Are Modulated by Second Sphere Residues. PLOS Pathogens 10: e1003817.

González, L.J., Stival, C., Puzzolo, J.L., Moreno, D.M., and Vila, A.J. (2018) Shaping Substrate Selectivity in a Broad-Spectrum Metallo-β-Lactamase. Antimicrobial Agents and Chemotherapy 62: e02079–02017.

González, L.J., Bahr, G., Nakashige, T.G., Nolan, E.M., Bonomo, R.A., and Vila, A.J. (2016) Membrane anchoring stabilizes and favors secretion of New Delhi metallo-β-lactamase. Nature chemical biology 12: 516–522.

Høiby, N., and Frederiksen, B. (2000) Microbiology of cystic fibrosis. In Cystic Fibrosis. London: Arnold Publishers - International book and journal publishers pp. 83–107.

Jacoby, G.A. (2009) AmpC β-lactamases. Clinical microbiology reviews 22: 161–182.

Lahiri, S.D., Johnstone, M.R., Ross, P.L., McLaughlin, R.E., Olivier, N.B., and Alm, R.A. (2014) Avibactam and class C beta-lactamases: mechanism of inhibition, conservation of the binding pocket, and implications for resistance. Antimicrob Agents Chemother 58: 5704–5713.

Lahiri, S.D., Walkup, G.K., Whiteaker, J.D., Palmer, T., McCormack, K., Tanudra, M.A. et al. (2015) Selection and molecular characterization of ceftazidime/avibactam-resistant mutants in *Pseudomonas aeruginosa* strains containing derepressed AmpC. J Antimicrob Chemother 70: 1650–1658.

Lister, P.D., Wolter, D.J., and Hanson, N.D. (2009) Antibacterial-resistant *Pseudomonas aeruginosa*: clinical impact and complex regulation of chromosomally encoded resistance mechanisms. Clin Microbiol Rev 22: 582–610.

Lujan, A.M., Moyano, A.J., Segura, I., Argarana, C.E., and Smania, A.M. (2007) Quorum-sensing-deficient (*lasR*) mutants emerge at high frequency from a *Pseudomonas aeruginosa mutS* strain. Microbiology 153: 225–237.

Luján, A.M., Maciá, M.D., Yang, L., Molin, S., Oliver, A., and Smania, A.M. (2011) Evolution and adaptation in *Pseudomonas aeruginosa* biofilms driven by mismatch repair system-deficient mutators. PLOS ONE 6: e27842.

Macia, M.D., Blanquer, D., Togores, B., Sauleda, J., Perez, J.L., and Oliver, A. (2005) Hypermutation is a key factor in development of multiple-antimicrobial resistance in *Pseudomonas aeruginosa* strains causing chronic lung infections. Antimicrob Agents Chemother 49: 3382–3386.

Mack, A.R., Barnes, M.D., Taracila, M.A., Hujer, A.M., Hujer, K.M., Cabot, G. et al. (2020) A Standard Numbering Scheme for Class C β-Lactamases. Antimicrobial Agents and Chemotherapy 64: e01841–01819.

MacLean, R.C., Hall, A.R., Perron, G.G., and Buckling, A. (2010) The population genetics of antibiotic resistance: integrating molecular mechanisms and treatment contexts. Nature Reviews Genetics 11: 405–414.

MacVane, S.H., Pandey, R., Steed, L.L., Kreiswirth, B.N., and Chen, L. (2017) Emergence of ceftolozane-tazobactam-resistant *Pseudomonas aeruginosa* during treatment is mediated by a single AmpC structural mutation. Antimicrob Agents Chemother 61.

Marvig, R.L., Johansen, H.K., Molin, S., and Jelsbak, L. (2013) Genome analysis of a transmissible lineage of *Pseudomonas aeruginosa* reveals pathoadaptive mutations and distinct evolutionary paths of hypermutators. PLOS Genetics 9: e1003741.

Matic, I. (2019) Mutation Rate Heterogeneity Increases Odds of Survival in Unpredictable Environments. Molecular Cell 75: 421–425.

Mehlhoff, J.D., Stearns, F.W., Rohm, D., Wang, B., Tsou, E.-Y., Dutta, N. et al. (2020) Collateral fitness effects of mutations. Proceedings of the National Academy of Sciences 117: 11597–11607.

Meini, M.-R., Tomatis, P.E., Weinreich, D.M., and Vila, A.J. (2015) Quantitative Description of a Protein Fitness Landscape Based on Molecular Features. Molecular Biology and Evolution 32: 1774–1787.

Mena, A., Smith, E.E., Burns, J.L., Speert, D.P., Moskowitz, S.M., Perez, J.L., and Oliver, A. (2008) Genetic adaptation of *Pseudomonas aeruginosa* to the airways of cystic fibrosis patients is catalyzed by hypermutation. J Bacteriol 190: 7910–7917.

Montanari, S., Oliver, A., Salerno, P., Mena, A., Bertoni, G., Tummler, B. et al. (2007) Biological cost of hypermutation in *Pseudomonas aeruginosa* strains from patients with cystic fibrosis. Microbiology 153: 1445–1454.

Morán-Barrio, J., Lisa, M.-N., Larrieux, N., Drusin, S.I., Viale, A.M., Moreno, D.M. et al. (2016) Crystal Structure of the Metallo-β-Lactamase GOB in the Periplasmic Dizinc Form Reveals an Unusual Metal Site. Antimicrobial Agents and Chemotherapy 60: 6013–6022.

Moya, B., Dotsch, A., Juan, C., Blazquez, J., Zamorano, L., Haussler, S., and Oliver, A. (2009) Beta-lactam resistance response triggered by inactivation of a nonessential penicillin-binding protein. PLoS Pathog 5: e1000353.

Moyano, A.J., Lujan, A.M., Argarana, C.E., and Smania, A.M. (2007) MutS deficiency and activity of the error-prone DNA polymerase IV are crucial for determining *mucA* as the main target for mucoid conversion in *Pseudomonas aeruginosa*. Mol Microbiol 64: 547–559.

Munita, J.M., Aitken, S.L., Miller, W.R., Perez, F., Rosa, R., Shimose, L.A. et al. (2017) Multicenter evaluation of ceftolozane/tazobactam for serious infections caused by carbapenem-resistant *Pseudomonas aeruginosa*. Clinical Infectious Diseases 65: 158–161.

Oliver, A. (2020) Antibiotic Resistance and Pathogenicity of Bacterial Infections Group - IdISBa.

Oliver, A., Canton, R., Campo, P., Baquero, F., and Blazquez, J. (2000) High frequency of hypermutable *Pseudomonas aeruginosa* in cystic fibrosis lung infection. Science 288: 1251–1254.

Palmer, A.C., and Kishony, R. (2013) Understanding, predicting and manipulating the genotypic evolution of antibiotic resistance. Nature reviews Genetics 14: 243–248.

Powers, R.A., Caselli, E., Focia, P.J., Prati, F., and Shoichet, B.K. (2001) Structures of Ceftazidime and Its Transition-State Analogue in Complex with AmpC β-Lactamase: Implications for Resistance Mutations and Inhibitor Design. Biochemistry 40: 9207–9214.

Prickett, M.H., Hauser, A.R., McColley, S.A., Cullina, J., Potter, E., Powers, C., and Jain, M. (2017) Aminoglycoside resistance of *Pseudomonas aeruginosa* in cystic fibrosis results from convergent evolution in the *mexZ* gene. Thorax 72: 40–47.

Raimondi, A., Sisto, F., and Nikaido, H. (2001) Mutation in *Serratia marcescens* AmpC β-Lactamase Producing High-Level Resistance to Ceftazidime and Cefpirome. Antimicrobial Agents and Chemotherapy 45: 2331–2339.

Rodríguez-Martínez, J.-M., Poirel, L., and Nordmann, P. (2009) Extended-Spectrum Cephalosporinases in *Pseudomonas aeruginosa*. Antimicrobial Agents and Chemotherapy 53: 1766–1771.

Rodriguez-Martinez, J.M., Poirel, L., and Nordmann, P. (2009) Molecular epidemiology and mechanisms of carbapenem resistance in *Pseudomonas aeruginosa*. Antimicrob Agents Chemother 53: 4783–4788.

Stiffler, Michael A., Hekstra, Doeke R., and Ranganathan, R. (2015) Evolvability as a Function of Purifying Selection in TEM-1 β-Lactamase. Cell 160: 882–892.

Thomas, V.L., McReynolds, A.C., and Shoichet, B.K. (2010) Structural Bases for Stability–Function Tradeoffs in Antibiotic Resistance. Journal of Molecular Biology 396: 47–59.

Tsutsumi, Y., Tomita, H., and Tanimoto, K. (2013) Identification of novel genes responsible for overexpression of *ampC* in *Pseudomonas aeruginosa* PAO1. Antimicrob Agents Chemother 57: 5987–5993.

Weinreich, D.M., Delaney, N.F., DePristo, M.A., and Hartl, D.L. (2006) Darwinian Evolution Can Follow Only Very Few Mutational Paths to Fitter Proteins. Science 312: 111–114.

Windels, E.M., Van den Bergh, B., and Michiels, J. (2020) Bacteria under antibiotic attack: Different strategies for evolutionary adaptation. PLOS Pathogens 16: e1008431.

Zasowski, E.J., Rybak, J.M., and Rybak, M.J. (2015) The β-Lactams Strike Back: Ceftazidime-Avibactam. Pharmacotherapy: The Journal of Human Pharmacology and Drug Therapy 35: 755–770.

