## Supplementary material for "Development of antibiotic resistance reveals diverse evolutionary pathways to face the complex and dynamic environment of a long-term treated patient": SI Appendix

### Supplementary Information for

#### Evolution of collateral resistance by b-lactamase remodeling is fueled by hypermutator *Pseudomonas aeruginosa* upon long-term antibiotic therapy

Claudia A. Colque[1,2], Pablo E. Tomatis[3,4¶], Andrea G. Albarracín Orio[1,2,5¶], Gina Dotta[3], Diego M. Moreno[6], Laura G. Hedemann[1,2], Rachel A. Hickman[7,8], Lea M. Sommer[7,8], Sofía Feliziani[1,2], Alejandro J. Moyano[1,2], Robert A. Bonomo[9,10], Helle K. Johansen[7,8,11], Søren Molin[8], Alejandro J. Vila[3,4*], Andrea M. Smania[1,2*]

[1]Universidad Nacional de Córdoba, Facultad de Ciencias Químicas, Departamento de Química Biológica Ranwel Caputto, Córdoba, Argentina.

[2]CONICET, Universidad Nacional de Córdoba, Centro de Investigaciones en Química Biológica de Córdoba (CIQUIBIC), Córdoba, Argentina.

[3] Instituto de Biología Molecular y Celular de Rosario (IBR), CONICET, Universidad Nacional de Rosario, Rosario, Argentina.

[4]Area Biofísica, Facultad de Ciencias Bioquímicas y Farmacéuticas, Universidad Nacional de Rosario, Argentina.

[5]IRNASUS, Universidad Católica de Córdoba, CONICET, Facultad de Ciencias Agropecuarias, Córdoba, Argentina

[6]IQUIR, Instituto de Química de Rosario, Universidad Nacional de Rosario, Santa Fe, Argentina

[7]Department of Clinical Microbiology, Rigshospitalet, Copenhagen, Denmark

[8]Novo Nordisk Foundation Centre for Biosustainability, Technical University of Denmark, Lyngby, Denmark

[9]Department of Molecular Biology and Microbiology, Case Western Reserve University, Cleveland, Ohio, USA

[10]Research Service, Louis Stokes Cleveland Department of Veterans Affairs, Cleveland, Ohio, USA

[11]Department of Clinical Medicine, University of Copenhagen, Copenhagen, Denmark

¶These authors contributed equally to this work.
*Correspondence to:
Andrea M. Smania, Centro de Investigaciones en Química Biológica de Córdoba (CIQUIBIC-CONICET), Universidad Nacional de Córdoba, X5000HUA, Córdoba, Argentina,.
Alejandro J. Vila, Universidad Nacional de Rosario, Instituto de Biología Molecular y Celular de Rosario (IBR-CONICET), S2000EZP, Rosario, Argentina,.

**This PDF file includes:**

Supplementary Materials and Methods
Tables supplement 1 to 4
Figures supplement 1 to 5
Supplementary Information References

#### Supplementary Materials and Methods

##### Data Availability Statement

The sequences of the *bla*$_{PDC}$ gene corresponding to the described PDCs variants have been deposited in GenBank at https://www.ncbi.nlm.nih.gov/genbank/ under the accession numbers shown in parentheses: BankIt2402117 PDC-458 (MW287261); BankIt2402134 PDC-459 (MW287262): BankIt2402138 PDC-460 (MW287263); BankIt2402139 PDC-461 (MW287264); BankIt2402140 PDC-462 (MW287265); BankIt2402141 PDC-463 (MW287266); BankIt2402142 PDC-464 (MW287267); BankIt2402144 PDC-465 (MW287268); BankIt2402146 PDC-466 (MW287269). Ultra-deep sequencing data is available under BioProject at https://www.ncbi.nlm.nih.gov/sra/PRJNA729779, accession number PRJNA729779.

##### Sequence analysis of *bla*$_{PDC}$ gene in *P. aeruginosa* CFD isolates

The *bla*$_{PDC}$ gene was amplified by PCR directly from bacterial colonies with primers *bla*$_{PDC}$_FOR and *bla*$_{PDC}$_REV (Table supplement 4). PCR amplifications were performed with the following conditions: 5min at 95°C, 30 cycles of 1 min at 95°C, 50 sec at 52°C, 2 min at 72°C, and a final extension of 10 min at 72°C. PCR products were cleaned with a Silica Purification Kit (Thermo Scientific), and sequenced directly using the same PCR primers (DNA Sequencing Facility, Univ. of Chicago, IL, USA). To score mutations within the gene, *bla*$_{PDC}$ sequences were compared with the corresponding gene sequence of the reference strain PAO1 (www.pseudomonas.com) using the CLC Genomics Workbench 10.1.1.

##### DNA extraction from sputum samples

For whole gene sequence analysis, genomic DNA was extracted directly from single sputum samples. For this purpose, sputum samples were thaw on ice, and 500 µL of sputum was treated with 30 µL (1M) Tris (2-carboxyethyl) phosphine, 10 µL proteinase K (20mg/mL) (Thermo Scientific) and 1 mL of DNA shield (Zymo Research) and vortexed for 30 sec. Samples were then added to 2 mL impact resistant screw-top tubes with 300 µL zirconia/glassbeads with a diameter of 0.1 mm (Carl Roth International) and vortexed on a secure horizontal holder at maximum speed for 5 min. Genomic DNA was extracted from the supernatant by ZR-Duet RNA/DNA mini-prep kit (Zymo Research) according to manufacturer's protocol.

**PCR amplification of $bla_{PDC}$ gene from whole sputum samples**

1.5µl of genomic DNA was used as template for gene amplification using Phusion High-Fidelity PCR master mix (ThermoFischer Scientific) with primers $bla_{PDC}$_FOR and $bla_{PDC}$_REV (Table supplement 4). The PCR amplification was performed as follows: 3 min at 98°C, 30 cycles of 10 sec at 98°C, 30 sec at 58°C, 2 min at 72°C and final extension of 5 min at 72°C. To validate correct amplification, amplicons were inspected by gel electrophoresis with GelRed (Biotium).

**Library preparation**

PCR product was cleaned with 1.8x AMPure beads (Agencourt®) and 15-30 ng of DNA was used as input for library preparation using KAPPA Hyper Plus kit (Kapa Biosystems) and barcoded using HT-Truseq dual-index adapter kit (Illumina Inc). Library was measured for DNA concentration with Qubit dsDNA HS Assay kit on the Qubit Fluorometer (ThermoFischer Scientific Inc 2015). The average size of the library was measured on the Bioanalyzer with DNA 1000 chip (Agilent Technologies). The sequencing library was sequenced on the MiSeq platform using the MiSeq V2 2x150bp read length kit (Illumina Inc).

**Sequence analysis**

Raw sequencing data were processed and trimmed to remove low quality reads. Trimmed reads were aligned to the gene sequence of the PAO1 reference strain (www.pseudomonas.com) by using the CLC Genomics Workbench 10.1.1 (Qiagen) and then both, forward and reverse files, were concatenated into a single file. SNPs were called by using the Low Frequency Variant detector package of CLC Workbench setting the following parameters: min frequency of the variant in the population of 2%, min count of 10 reads supporting the nucleotide position, and min count of 2 reads supporting the variation. From the output file, the only entries considered were the ones with an average quality above 25 and a forward/reverse balance around 0.5. Sequence variants were classified as synonymous (S) or non-synonymous (NS) and the frequency (%) in the population was scored.

**Construction of *P. aeruginosa* Δ*bla*$_{\text{PDC}}$ deficient strain (PAΔA)**

Strain was constructed by allelic replacement using pKNG101 vector (Kaniga et al., 1991) and primers *bla*$_{\text{PDC}}$_FOR_Up *bla*$_{\text{PDC}}$_REV_Up, *bla*$_{\text{PDC}}$_FOR_Down and *bla*$_{\text{PDC}}$_REV_Down (Table supplement 4). Briefly, amplicons of 467 and 260 bp from the outside regions of *bla*$_{\text{PDC}}$ generated with latter primers, were blunt-ended and then ligated to give a 727 bp fragment which was clone into ApaI and SpeI sites of pKNG101 vector. Conjugation experiments were performed by biparental mating of *E. coli* SM10 pKNG:727 and PAO1. Positive clones were selected by the ability to grow on sucrose 25% and sensibility to streptomycin 50 μg/mL. Phenotypic analysis to confirm the deletion of *bla*$_{\text{PDC}}$ in the mutant was analyzed by growth inhibition on cefoxitin disks (FOX30μg Britania).

**Construction of *P. aeruginosa* *bla*$_{\text{PDC}}$-*lacZ* strain (PAΔA-*lacZ*)**

The PAΔA-*lacZ* strain was constructed by transformation of PAΔA strain with the pUC18-mini Tn7T and pTNS1 plasmids (Table supplement 4) by insertion of the β-galactosidase gene at the single attTn7 site downstream of the *glmS* gene in the *P. aeruginosa* genome (Choi et al., 2005).

In order to obtain an unmarked PAΔA-*lacZ* strain, the gentamicin cassette was removed by transforming with pFLP2 vector (Table supplement 4). Positive clones were selected for the ability to grow on sucrose 10% and sensibility to gentamicin 40 μg/mL and carbenicillin 200 μg/mL. Expression of β-galactosidase (blue colonies) was visualized by the addition of 100 μg/mL X-gal.

**Cloning of *bla*$_{\text{PDC}}$ allelic variants**

*bla*$_{\text{PDC}}$ allelic variants obtained from the CFD collection were cloned into pMBLe (González et al., 2016). For this purpose, the complete gene sequence (including native peptide leader) of *bla*$_{\text{PDC}}$ was amplified by PCR, from bacterial colonies, with primers *bla*$_{\text{PDC}}$_FOR_NdeI and *bla*$_{\text{PDC}}$_REV_HindIII (for susceptibility testing) and with *bla*$_{\text{PDC}}$_FOR_NdeI and *bla*$_{\text{PDC}}$_REV_HindIII_ST (for Western Blot analyses) (Table supplement 4), cloned into NdeI and HindIII sites in pMBLe and transformed into *Escherichia coli* DH5α chemically competent by $CaCl_2$. Transformants were selected on LB agar supplemented with 10 μg/mL of gentamicin. After sequencing step to ensure that no mutation was introduce during PCR amplification, the resulting pMBLe plasmids (Table supplement 4) were transferred by electroporation (Bio-Rad MicroPulser) into the

PA PAΔA knockout mutant. Transformants were selected on gentamicin 40 μg/mL. The expression of *bla*$_{PDC}$ was induced by addition of 10 μM IPTG verifying that the PAΔA MIC transformed with pMBLe expressing PDC-1 was comparable to that of PAO1 (Table supplement 3). We also verified that the C-terminus Strep tag (added to the cloning for Western Blot analyses purposes), did not affect the ability of PDC to confer resistance.

**PDC expression levels in pMBLe**

The *bla*$_{PDC}$ expression levels from pMBLe induced with IPTG were evaluated by Western Blot assays. For this purpose, *bla*$_{PDC}$ allelic variants were labeled with a C-terminal Strep-tag. PAΔA strain was transformed with the pMBLe carrying the different variants, bacteria were grown ON in LB in the presence of 0, 10 and 25μM of IPTG supplemented with 40 μg/ml of gentamicin. Then, 1 mL of each culture was pelleted and resuspended in 20 mM Tris-HCl (pH 7.4), 0.5 M NaCl, 15% glycerol, 1 mM phenylmethylsulfonylfluoride and 1 mM benzamidine protease inhibitors. 25 μg of total proteins (measure with Bradford) was separated through sodium dodecyl sulfate (SDS)- polyacrylamide gel electrophoresis (PAGE) 10%, then the proteins were transferred to nitrocellulose membranes (0.22 μm, Sigma) for 1 hour at 300 mA. The blots were blocked for one hour in a 5% milk in phosphate-buffered saline (PBS) solution at room temperature. Incubation with primary antibody (mouse anti-Strep II monoclonal, IBA) was added at 1/10,000 overnight at 4˚C in 5% milk/PBS, then washings were performed with PBS/Tween 20, and the secondary antibody (IR-Dye 800 anti-mouse, LI-COR Bioscience) was added at a 1:10,000 dilution for 1 hour in 5% milk/PBS. Membranes were scanned on the Odyssey infrared imager instrument (LI-COR Bioscience).

**Drug susceptibility testing**

MIC determinations were performed by broth dilution method according to CLSI guidelines (CLSI, 2019). The β-lactam antibiotics tested (breakpoints shown as ≤susceptible/≥resistant) were: ceftazidime (8/32 μg/mL), cefepime (8/32 μg/mL), ceftolozane (4/16 μg/mL), ceftolozane/tazobactam (4-4/16-4 μg/mL), piperacillin (16/128 μg/mL), piperacillin/tazobactam (16-4/128-4 μg/mL), aztreonam (8/32 μg/mL), imipenem (2/8 μg/mL) and meropenem (2/8 μg/mL). *P. aeruginosa* ATCC 27853 was used as control strain.

**Competition experiments**

Competitive fitness of *bla*$_{PDC}$ variants was determined by direct competition of each variant PDC-461, PDC-462, PDC-463 and PDC-464 with their ancestral *bla*$_{PDC}$ PDC-3 as well as among variants. For this purpose, PAΔA and PAΔA-*lacZ* strains were transformed with pMBLe-PDC-3, pMBLe-PDC-461, pMBLe-PDC-462, pMBLe-PDC-463 and pMBLe-PDC-464 plasmids to obtain PAΔA-3, PAΔA-461, PAΔA-462, PAΔA-463 and PAΔA-464 strains. For competition experiments, co-cultures at a ratio of 1:1 were inoculated to a final OD of 0.1 and grown in LB medium supplemented with 40 µg/mL gentamicin, 10 µM IPTG, and the presence or absence of different sub_MIC concentrations of ceftazidime or aztreonam for 24 h at 37˚C with shaking. The concentration of antibiotic for a specific competition was defined according to the MIC of the lesser advantage variant. For ceftazidime the concentrations used were as follows: 4xMIC of PDC-3 (16 µg/mL) for competitions relative to PAΔA-3; 1/2xMIC of PDC-464 (32 µg/mL) for competitions relative to PAΔA-464; 1/2xMIC of PDC-463 (32 µg/mL) and 1/4MIC of PDC-462 (32 µg/mL) when competed each other, considering for the latter that 32µg/mL would be enough to differ between PAΔA-462 and PAΔA-461 (MICs of 128 µg/mL). For aztreonam the concentrations used for each experiment were: a 4xMIC of PDC-3 (16 µg/mL) for competitions relative to PAΔA-3; 1/2MIC of PDC-464 (8 µg/mL) for competitions relative to PAΔA-464; 1/2MIC of PDC-461 (16 µg/mL) for competitions relative to PAΔA-461 and 1/2MIC of PDC-462 (32 µg/mL) when competed against PAΔA-462.

Afterwards, 100 to 300 cells from the final culture were plated on LB agar plates supplemented with 100 µg/mL X-Gal and grown for 16 h at 37˚C. Blue-white colony screening was used to determine the proportion of *lacZ*+ and *lacZ*-, and thus of each variant in the population after competition. To calculate fitness (S) the following equation was used:

$$S = \ln\left(\frac{\delta f}{\delta i}\right) \mathrm{a} - \ln\left(\frac{\delta f}{\delta i}\right) \mathrm{b}$$

Where $\delta i$ and $\delta f$ are the number of CFU/mL of initial and final co-cultures; *a* and *b* represent the two competing strains. When S=0, both strains compete the same, and when S>0 meaning that strain *a* out-competes strain *b*. Two independent experiments, each with three replicates, were assessed for each competition. Two-way analysis of variance (ANOVA) followed by Tukey's Multiple Comparisons Test of relative fitness values were performed using GraphPad Prism 7.0 Software. Statistically significant differences ($P < 0.05$) were recorded.

**Expression and purification of PDC proteins**

Each *bla*$_{PDC}$ gene encoding mature versions, residues 27 to 397 of the full-length protein, were PCR amplified with primers *bla*$_{PDC}$_FOR_Mature and *bla*$_{PDC}$_REV_HindIII (Table supplement 4). Then, mature versions were cloned into NdeI and HindIII sites as N-terminal fusion to a 6xHis-tag in the expression vector pET28bTEV (González et al., 2016), and transformed into *E. coli* DH5α. Transformants were selected in LB agar plates supplemented with 25 μg/mL of kanamycin. Each pET-PDC plasmid was isolated, verified by PCR and sequencing, and then used to transform *E. coli* BL21 (DE3) strain. For PDC overexpression the strain (BL21::pET-PDC variant) was grown with 25 μg/mL of kanamycin at 37 °C in one liter of LB medium until it reached $OD_{600nm}$ of 0.6, when protein expression was induced by addition of 0.5 mM IPTG, following an incubation with agitation at 20˚C for 20 h. Cell were harvested by centrifugation, resuspended in 25 mL of 50 mM Tris and 200 mM NaCl (Buffer A, pH 8.0) supplemented with DNAse (10 μg/ml) and 50 mM $MgCl_2$. Then, cells were disrupted by sonication (5 times at 40% for 30s and 5 min pause). Afterwards, lysed cells were centrifuged for 1 h at 16000 rpm to remove insoluble materials. Supernatant was loaded at 2 mL/min on a 5 mL HisTrap HP column previously equilibrated with buffer A at 4˚C. Bounded protein to the Ni-Sepharose resin was eluted with a lineal gradient of Buffer A + 500mM Imidazol (pH 8.0). Active fractions were pooled and concentrated using Amicon Ultra 10-15 K down to 5 mL and then subjected to dialysis, for elimination of Imidazol, with 25 mm cellulose membrane against Buffer A at 4 ˚C for 16 h. His-PDC was then digested with His-tagged TEV protease in 30:1 ratio for 2 h at room temperature. PDC protein was then loaded on the HisTrap column and eluted with Buffer A. Purified β-lactamases were concentrated by ultrafiltration using Amicon Ultra 10-15 K to a final concentration of 10 to 30 mg/mL and store at -20˚C. PDC mature protein concentrations were determined from the absorbance at 280 nm using a molar absorption coefficient $\varepsilon_{280}$ of 55,800 $M^{-1}$ $cm^{-1}$ (calculated using Expasy ProtParam, available at http:// web.expasy.org/protparam/). All final protein preparations have a purity > 95%, as determined by SDS-PAGE.

**Steady-state kinetic measurements**

Purified β-lactamases PDC-3, PDC-461, PDC-462 and PDC-463 were used to determine kinetic parameters using ceftazidime ($\Delta\varepsilon_M^{260nm}$ = 9,000 $M^{-1}$ $cm^{-1}$), piperacillin ($\Delta\varepsilon_M^{235nm}$ = 820 $M^{-1}$ $cm^{-1}$) and imipenem ($\Delta\varepsilon_M^{300nm}$ = 9,000 $M^{-1}$ $cm^{-1}$), and ceftolozane ($\Delta\varepsilon_M^{260nm}$ = 8,000 $M^{-1}$ $cm^{-1}$) as substrates. The initial reaction rates at different substrate concentrations were analyze with a Jasco V-670 spectrophotometer at 30˚C in 10 mM phosphate buffer (pH 7.0) in a 0.1cm or 1cm cuvette when appropriate. Dependences of initial rates on substrate concentration were analyzed by a nonlinear least squares fit of the data with Michaelis-Menten equation using GraphPad Prism 7.0 in order to determine $K_M$ and $k_{cat}$ values. Reported kinetic parameters correspond to averages from at least two determinations with independent protein samples.

**Molecular modeling**

**(i) Initial structures.** The initial structure of PDC-3, was built *in-silico* replacing Thr by Ala in position 79 from the crystallographic structure of *Pseudomonas aeruginosa* class C beta-lactamase PDC-1 obtained from the Protein Data Bank entry 4OOY (Lahiri et al., 2014). The mutants PDC-461, PDC-462 and PDC-463 were constructed replacing the corresponding amino acids.

**(ii) Classical molecular dynamic simulations**

MD simulations were performed starting from initial structures built *in-silico* as described above. Each protein was immersed in a truncated octahedral periodic box with a minimum solute-wall distance of 8 Å, filled with explicit TIP3P water molecules (Jorgensen et al., 1983) using the AMBER16 leap module (D.A. Case et al., 2016). Molecular dynamic simulations were performed with the AMBER16 package (D.A. Case et al., 2016), using the ff14SB (Maier et al., 2015) force field. Particle-mesh Ewald (PME) was implemented for long range interactions with a cutoff distance of 12 Å (Luty et al., 1995). Temperature and pressure were regulated with the Berendsen thermostat and barostat, as implemented in the AMBER16 (D.A. Case et al., 2016), using a time constant of 2 ps (Berendsen et al., 1984). All bonds involving hydrogen were fixed using the SHAKE algorithm (Ryckaert et al., 1977). Each initial system was minimized using a multistep protocol, then heated from 0 to 300 K, and finally a short simulation at constant temperature of 300 K, under constant pressure of 1 bar, was performed to allow the systems to reach proper density. These equilibrated structures were the starting point for 200 ns of MD simulations at 300 K in the NVT ensemble. This protocol was used previously (González et al., 2014; Morán-Barrio et al., 2016; González et al., 2018).

To analyze the MD simulations, different parameters (Root Mean Square Deviation, Root Mean Square Fluctuation, distances, etc.) were obtained with the cpptraj module (Roe and Cheatham, 2013). Conformational clusterization was performed using the hierarchical agglomerative approach from the cpptraj module of Amber16 (Roe and Cheatham, 2013).

**(iii) QM-MM calculations**

For hybrid QM–MM calculations, we used Self-Consistent Charge Density Functional Tight Binding (SCC-DFTB) (Gaus et al., 2011) to describe the QM region as implemented in Amber16 and the same force field used in the classical MD simulations to describe the MM region (Seabra et al., 2007; D.A. Case et al., 2016). We extracted representative structures of each variant from the MD simulations and performed an structural alignment of them using the VMD software (Humphrey et al., 1996) with the crystallographic structure of a substrate bound PDC (Protein Data Bank entry 1IEL) (Powers et al., 2001). The C-N bond of the beta-lactam ring of the ceftazidime was rebuilt *in silico* to obtain an initial structure of a protein-ceftazidime complex. The simulation protocol consists of an initial minimization at the molecular mechanic level of each complex structure to accommodate solvent molecules and possible clashes, followed by QM-MM geometry optimization. The QM region consisted of the residue of Ser64 and the ceftazidime. We perform two approaches, one with a geometry optimization without restrictions and another with a distance restraint of 2.2 Å between the oxygen atom of Ser64 and the C atom of the carbonyl group of the ceftazidime. We also performed the QM-MM calculations in two steps, first we applied a distance restraint of 2.2 Å between the oxygen atom of Ser64 and the C atom of the carbonyl group of the ceftazidime to accommodate the substrate in catalytic conformation and then we removed the restraint and a full geometry optimization was done.

#### Supplementary Tables

**Table supplement 1.** Non synonymous mutations found within CFD isolates and their representation throughout PDC database

| DNA change | AA change [a] | Number of PDCs [b] (%) | PDC variant [c] | References [d] |
|---|---|---|---|---|
| A313G | T79A | 413 (90.4%) | PDC-3 | (Rodríguez-Martínez et al., 2009) |
| C344T | A89V | 0 (0%) | PDC-460 to PDC-465 | This study |
| C436A | Q120K | 6 (1.3%) | PDC-459, PDC-461 to PDC-464, PDC-466 | This study |
| C539T | P154L | 10 (2.2%) | PDC-73, PDC-459 | (Berrazeg et al., 2015) |
| C643T | H189Y | 12 (2.6%) | PDC-463 and PDC-464 | (Marvig et al., 2013; Haidar et al., 2017), this study |
| G692A | G205D | 0 (0%) | PDC-466 | This study |
| T716C | V213A | 50 (10.9%) | PDC-50, PDC-459 to -464, PDC-466 | (Berrazeg et al., 2015; López-Causapé et al., 2017) |
| A884C | T256P | 0 (0%) | PDC-465 | This Study |
| G724A | G216S | 6 (1.3%) | PDC-458 and PDC-465 | This study |
| A1040G | N321S | 21 (4.6%) | PDC-460, PDC-462, PDC-464 | This study |
| G1066A | V330I | 96 (21.0%) | PDC-458, PDC-7, PDC-75, PDC-86, PDC-87, PDC-88, PDC-90, PDC-91 | (Marvig et al., 2013; Berrazeg et al., 2015) |
| A1118T | N347I | 11 (2.4%) | PDC-76, PDC-84 | (Berrazeg et al., 2015) |

[a] Amino acid (AA) variations were defined according to the mature protein from PAO1 strain, after cleavage of the 26 N-terminal amino acid residues from the signal peptide. [b] Preponderance of the AA variation along the *Pseudomonas*-derived cephalosporinase

(PDC) database (Oliver, 2020) (updated November 17th 2020). Number indicates the number of PDCs carrying each AA variation and percentage out of the 457 PDCs is shown in parenthesis. [c] Simple and/or multiple PDC mutants harboring each mutation are included. Numbers of PDCs which have only been characterized before are specified. [d] Cited references include those mutations that have been characterized and/or described in CF *P. aeruginosa* isolates.

**Table supplement 2.** Results from Amplicon Sequencing with Illumina MiSeq

| Reference position [A] | Count | Coverage | Frequency | For/rev balance | Average quality | Coding region change [b] | Amino acid change [c] |
|---|---|---|---|---|---|---|---|
| 316 | 86137 | 88089 | 97.78406 | 0.495699 | 36.34678 | 297C>A | |
| 332 | 87769 | 88210 | 99.50006 | 0.498832 | 36.28925 | 313A>G | Thr79Ala |
| 363 | 48489 | 88661 | 54.69034 | 0.494978 | 35.90165 | 344C>T | Ala89Val |
| 382 | 1939 | 88410 | 2.193191 | 0.48066 | 33.92264 | 363A>G | |
| 455 | 53187 | 85856 | 61.94908 | 0.498073 | 35.86974 | 436C>A | Gln120Lys |
| 616 | 94383 | 96793 | 97.51015 | 0.496784 | 36.66562 | 597G>A | |
| 662 | 3488 | 96707 | 3.606771 | 0.495413 | 37.00201 | 643C>T | His189Tyr |
| 709 | 93237 | 93570 | 99.64412 | 0.498665 | 36.67504 | 690T>C | |
| 727 | 93390 | 98579 | 94.7362 | 0.499946 | 36.15308 | 708G>C | |
| 735 | 74750 | 93001 | 90.37548 | 0.49996 | 35.4617 | 716T>C | Val213Ala |
| 1059 | 46643 | 93087 | 50.10689 | 0.477049 | 36.91551 | 1040A>G | Asn321Ser |
| 1090 | 2468 | 89992 | 2.742466 | 0.462723 | 36.84643 | 1071G>A | |
| 1105 | 79771 | 81828 | 97.48619 | 0.467676 | 37.30258 | 1086C>A | |
| 1137 | 1586 | 69192 | 2.292173 | 0.465322 | 36.50631 | 1118A>T | Asn347Ile |

[a] Reads from Illumina were mapped against *ampC* gene from PAO1 reference strain (*www.pseudomonas.com*). [b] DNA changes C297A, A313G (T79A), G597A, T690C, G708C, C1086A appearing at around 98% of total reads are also present in all single isolates sequenced from CFD patient, and considered as common polymorphisms in CFD lineage. [c] Position refers to mature AmpC protein, after cleavage of the 26 N-terminal amino acid (AA) from signal peptide.

**Table supplement 3.** Ceftazidime MICs of PAΔA expressing AmpC variants in IPTG-inducible pMBLe

| | IPTG (μM) | 0 | 10 | 25 |
|---|---|---|---|---|
| | | CAZ MIC (μg/mL) [a] | | |
| Strains | PAΔA | 2 | ND | ND |
| | PAΔA-EV | 1 | 1 | 1 |
| | PAΔA-1 | 4 | 4 | 4 |
| | PAΔA-458 | 4 | 4 | 8 |

[a] For ceftazidime (CAZ) MIC determinations the agar diffusion method was performed according to CLSI guidelines. ND, not determined. MH medium, Gm 40 μg/mL was used with the corresponding concentration of CAZ and IPTG.

**Table supplement 4.** Strains, plasmids and primers used in this study

| Bacterial strains, plasmids or primers | Relevant properties or sequences (5’→ 3’) | Source or reference |
|---|---|---|
| Strains | | |
| PAO1 | Wild-type *P. aeruginosa* first isolated from hydrocarbon soils | (Holloway, 1955) |
| PAΔA | PDC-deficient mutant from PAO1 | This study |
| PAΔA-*lacZ* | PDC-deficient mutant from PAO1 expressing β-galactosidase | This study |
| *E. coli* DH5α | Laboratory *E. coli* strain | Invitrogen |
| *E. coli* SY327 | λpir recipient strain for suicide vector | (Miller and Mekalanos, 1988) |
| *E. coli* SM10 | λpir recipient strain for suicide vector and for conjugation | (Simon et al., 1983) |
| *E. coli* BL21 (DE3) | Protein expression | Novagen |
| Plasmids | | |
| pKNG | $Sm^R$, suicide vector derivative of Tn5 | (Kaniga et al., 1991) |
| pMBLe | $Gm^R$, *Nde*I and *Hind*III sites, Ptac promoter | (González et al., 2016) |
| pTNS1 | $Ap^R$, Helper plasmid DNA | (Choi et al., 2005) |
| pUC18-mini-Tn7T | $Gm^R$, lacZ transcriptional fusion vector | (Choi and Schweizer, 2006) |
| pFLP2 | $Amp^R$, for site-specific excision of gentamicin marker | (Hoang et al., 1998) |
| pET28bTEV | $Km^R$, expression vector N-terminally 6xHis-tagged proteins with TEV site replacing the thrombin cleavage site separating the His tag from cloned gene | (Tomalka et al., 2013) |
| pMBLe-PDC-1 | pMBLe vector expressing *ampC* allele from PAO1 | This study |
| pMBLe-PDC-3 | pMBLe vector expressing ampC allele from 1991 (T79A) | This study |
| pMBLe-PDC-458 | G216S, V330I | This study |
| pMBLe-PDC-459 | pMBLe vector expressing ampC allele with Q120K, P154L, V213A mutations | This study |
| pMBLe-PDC-460 | pMBLe vector expressing ampC allele with A89V, V213A, N321S mutations | This study |
| pMBLe-PDC-461 | pMBLe vector expressing ampC allele with A89V, Q120K, V213A mutations | This study |
| pMBLe-PDC-462 | pMBLe vector expressing ampC allele with A89V, Q120K, V213A, N321S mutations | This study |

| | | |
|---|---|---|
| pMBLe-PDC-463 | pMBLe vector expressing ampC allele with A89V, Q120K, H189Y, V213A mutations | This study |
| pMBLe-PDC-464 | pMBLe vector expressing ampC allele with A89V, Q120K, H189Y, V213A, N321S mutations | This study |
| Primers | | |
| $bla_{PDC}$_FOR_Up | 5’-TACAAAAAAGCAGGCTCGACCGTGAAGGTTCCGA-3’ | This study |
| $bla_{PDC}$_REV_Up | 5’-TCAGAGCGCTTTTGAAGCTAATTCGGAGGATTGGCGTCCTTTGT-3’ | This study |
| $bla_{PDC}$_FOR_Down | 5’-AGGAACTTCAAGATCCCCAATTCGGAGGGCGACGGAGCGTA-3’ | This study |
| $bla_{PDC}$_REV_Down | 5’-TACAAGAAAGCTGGGTATGCCCAGTTGTTCCTCCA-3’ | This study |
| $bla_{PDC}$_FOR_NdeI | 5’-GTCATATGCGCGATACCAGATT-3’ | This study |
| $bla_{PDC}$_REV_HindIII | 5’- ATAAGCTTTCAGCGCTTCAGCGGCA-3’ | This study |
| $bla_{PDC}$_REV_HindIII_ST | 5’-ATAAGCTTTCACTTTTCGAATTGTGGGTGAGACCAGCGCTTCAGCGGCA-3’ | This study |
| $bla_{PDC}$_FOR | 5’-ATCGCCGCTTCCACACTGCT-3’ | This study |
| $bla_{PDC}$_REV | 5’-CTGAGGATGGCGTAGGCGATCT-3’ | This study |
| $bla_{PDC}$_FOR_Mature | 5’ GCAGACATCATATGGGCGAGGCCCCGGCGGATC-3’ | This study |

$Sm^R$, streptomycin resistance, $Ap^R$, ampicillin resistance; $Gm^R$, gentamicin resistance, $Km^R$, kanamycin resistance

### Supplementary Figures

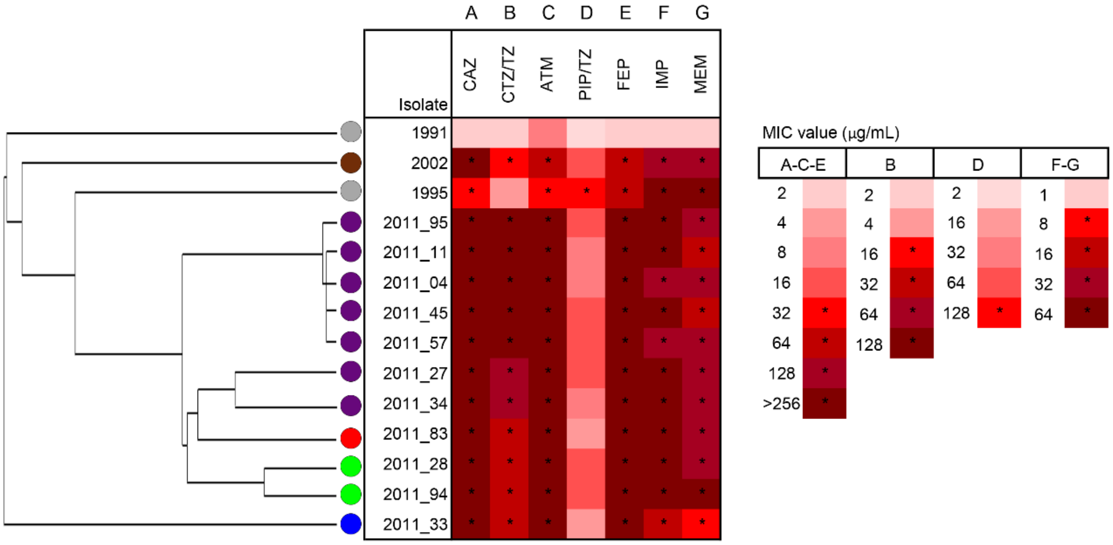


**Figure supplement 1. β-lactam antibiotic resistance profiles of *P. aeruginosa* isolates from the CFD lineage.**

Each column represents the MIC values of the different antibiotics tested: ceftazidime (CAZ), ceftolozane-tazobactam (CTZ/TZ), aztreonam (ATM), piperacillin-tazobactam (PIP/TZ), cefepime (FEP), imipenem (IMP) and meropenem (MEM). The red intensity indicates the MIC levels for each antibiotic. Asterisks indicate resistance according to CLSI guidelines. CAZ, ATM, TZP, FEP, IMP and MEM values are from Colque *et al* (Colque et al., 2020). The phylogenetic tree on the left represents the genetic clustering of isolates (rows), based on the result of maximum-parsimony analysis, and was constructed based on the accumulation of new SNPs relative to the sequences of the ancestor from 1991 (Feliziani et al., 2014). Circle colors represent the types of PDC variants that harbored each of the CFD isolate sequenced (i.e.: grey: PDC-3, brown: PDC-458, purple: PDC-462, red: PDC-460, green: PDC-463, blue: PDC-459).

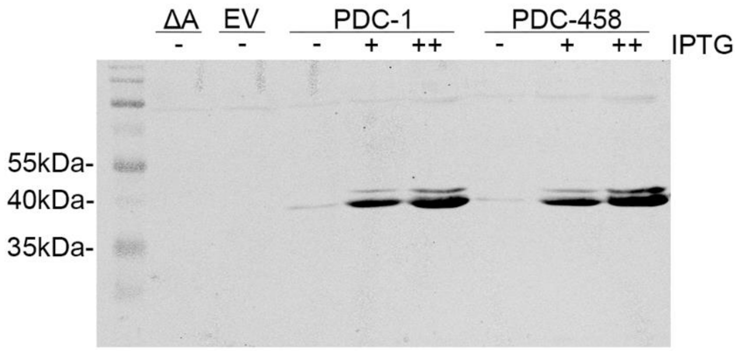


**Figure supplement 2. Representative Western Blot showing the controlled expression of PDC variants PDC-1 and PDC-458.**

Entire cell extracts of strain PAΔA expressing either strep-tagged PDC-1 or PDC-458 were used for Western blotting by using specific antibodies against strep-tag. Overnight bacterial cultures were induced with IPTG 10 μM (+), 20 μM (++), or grown without IPTG (-). Cell extracts of PAΔA transformed with the empty vector (EV) pMBLe or with no vector at all (ΔA) were used as negative controls.

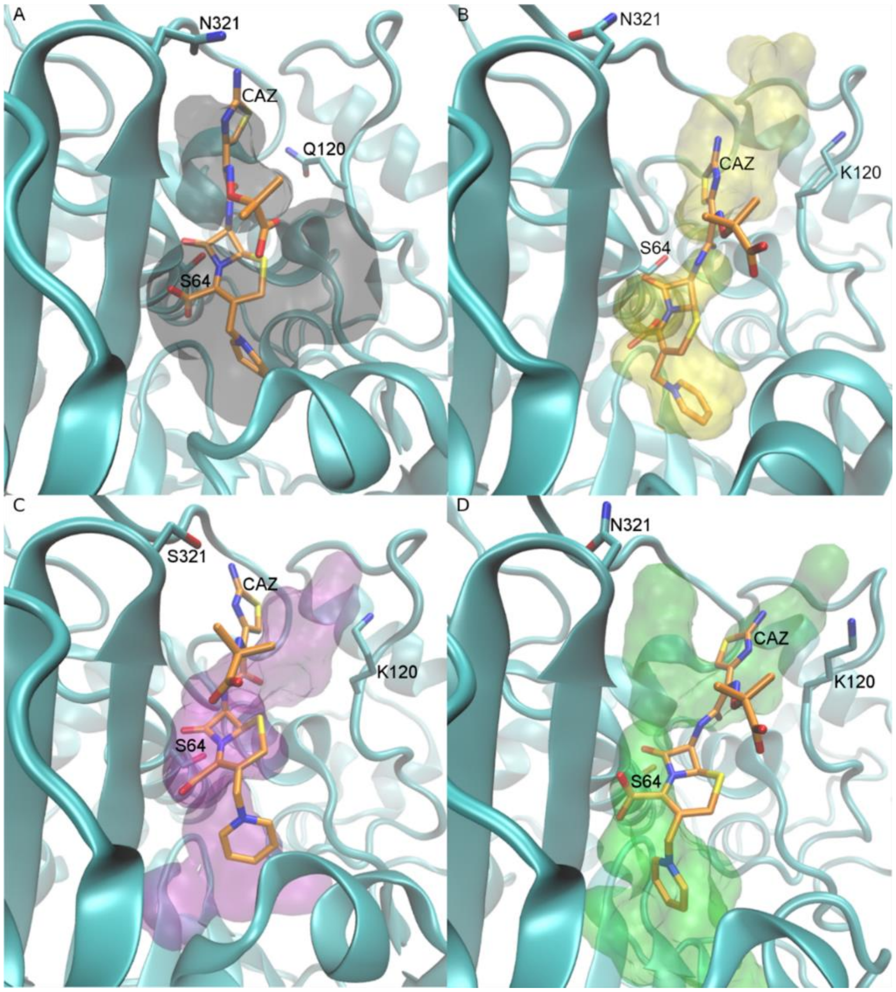


**Figure supplement 3. Representation of the active site volume cavity calculated in a representative snapshot of the MD simulations overlaid with the complex protein-ceftazidime optimized structures for each protein studied**.

N atoms are depicted in blue, O atoms in red, C atoms of the protein in cyan and C atom of the ceftazidime in orange, volume cavity is represented with a surface shape representation in different colors. A) PDC-3 (gray), B) PDC-461 (yellow), C) PDC-462 (purple) and D) PDC-463 (green).

#### SI References

Berendsen, H.J.C., Postma, J.P.M., Gunsteren, W.F.v., DiNola, A., and Haak, J.R. (1984) Molecular dynamics with coupling to an external bath. *The Journal of Chemical Physics* **81**: 3684-3690.

Berrazeg, M., Jeannot, K., Ntsogo Enguene, V.Y., Broutin, I., Loeffert, S., Fournier, D., and Plesiat, P. (2015) Mutations in beta-lactamase AmpC increase resistance of *Pseudomonas aeruginosa* isolates to antipseudomonal cephalosporins. *Antimicrob Agents Chemother* **59**: 6248-6255.

CLSI (2019) Performance Standards for Antimicrobial Susceptibility Testing. 29th ed. In: Wayne, PA: Clinical and Laboratory Standards Institute.

Colque, C.A., Albarracín Orio, A.G., Feliziani, S., Marvig, R.L., Tobares, A.R., Johansen, H.K. et al. (2020) Hypermutator *Pseudomonas aeruginosa* exploits multiple genetic pathways to develop multidrug resistance during long-term infections in the airways of cystic fibrosis patients. *Antimicrobial Agents and Chemotherapy*: AAC.02142-02119.

Choi, K.-H., and Schweizer, H.P. (2006) mini-Tn7 insertion in bacteria with single attTn7 sites: example *Pseudomonas aeruginosa*. *Nature Protocols* **1**: 153-161.

Choi, K.H., Gaynor, J.B., White, K.G., Lopez, C., Bosio, C.M., Karkhoff-Schweizer, R.R., and Schweizer, H.P. (2005) A Tn7-based broad-range bacterial cloning and expression system. *Nat Methods* **2**: 443-448.

D.A. Case, R.M. Betz, D.S. Cerutti, T.E. Cheatham, III, T.A. Darden et al. (2016) AMBER 2016. In: University of California, San Francisco.

Feliziani, S., Marvig, R.L., Luján, A.M., Moyano, A.J., Di Rienzo, J.A., Krogh Johansen, H. et al. (2014) Coexistence and within-host evolution of diversified lineages of hypermutable *Pseudomonas aeruginosa* in long-term cystic fibrosis infections. *PLOS Genetics* **10**: e1004651.

Gaus, M., Cui, Q., and Elstner, M. (2011) DFTB3: Extension of the Self-Consistent-Charge Density-Functional Tight-Binding Method (SCC-DFTB). *Journal of Chemical Theory and Computation* **7**: 931-948.

González, L.J., Moreno, D.M., Bonomo, R.A., and Vila, A.J. (2014) Host-Specific Enzyme-Substrate Interactions in SPM-1 Metallo-β-Lactamase Are Modulated by Second Sphere Residues. *PLOS Pathogens* **10**: e1003817.

González, L.J., Stival, C., Puzzolo, J.L., Moreno, D.M., and Vila, A.J. (2018) Shaping Substrate Selectivity in a Broad-Spectrum Metallo-β-Lactamase. *Antimicrobial Agents and Chemotherapy* **62**: e02079-02017.

González, L.J., Bahr, G., Nakashige, T.G., Nolan, E.M., Bonomo, R.A., and Vila, A.J. (2016) Membrane anchoring stabilizes and favors secretion of New Delhi metallo-β-lactamase. *Nature chemical biology* **12**: 516-522.

Haidar, G., Philips, N.J., Shields, R.K., Snyder, D., Cheng, S., Potoski, B.A. et al. (2017) Ceftolozane-Tazobactam for the Treatment of Multidrug-Resistant *Pseudomonas aeruginosa* Infections: Clinical Effectiveness and Evolution of Resistance. *Clinical Infectious Diseases* **65**: 110-120.

Hoang, T.T., Karkhoff-Schweizer, R.R., Kutchma, A.J., and Schweizer, H.P. (1998) A broad-host-range Flp-FRT recombination system for site-specific excision of chromosomally-located DNA sequences: application for isolation of unmarked *Pseudomonas aeruginosa* mutants. *Gene* **212**: 77-86.

Holloway, B.W. (1955) Genetic Recombination in *Pseudomonas aeruginosa*. *Microbiology* **13**: 572-581.

Humphrey, W., Dalke, A., and Schulten, K. (1996) VMD: Visual molecular dynamics. *Journal of Molecular Graphics* **14**: 33-38.

Jorgensen, W.L., Chandrasekhar, J., Madura, J.D., Impey, R.W., and Klein, M.L. (1983) Comparison of simple potential functions for simulating liquid water. *The Journal of Chemical Physics* **79**: 926-935.

Kaniga, K., Delor, I., and Cornelis, G.R. (1991) A wide-host-range suicide vector for improving reverse genetics in gram-negative bacteria: inactivation of the *blaA* gene of *Yersinia enterocolitica*. *Gene* **109**: 137-141.

Lahiri, S.D., Johnstone, M.R., Ross, P.L., McLaughlin, R.E., Olivier, N.B., and Alm, R.A. (2014) Avibactam and class C beta-lactamases: mechanism of inhibition, conservation of the binding pocket, and implications for resistance. *Antimicrob Agents Chemother* **58**: 5704-5713.

López-Causapé, C., Sommer, L.M., Cabot, G., Rubio, R., Ocampo-Sosa, A.A., Johansen, H.K. et al. (2017) Evolution of the *Pseudomonas aeruginosa* mutational resistome in an international cystic fibrosis clone. *Scientific reports* **7**: 5555-5555.

Luty, B.A., Tironi, I.G., and Gunsteren, W.F.v. (1995) Lattice-sum methods for calculating electrostatic interactions in molecular simulations. *The Journal of Chemical Physics* **103**: 3014-3021.

Maier, J.A., Martinez, C., Kasavajhala, K., Wickstrom, L., Hauser, K.E., and Simmerling, C. (2015) ff14SB: Improving the Accuracy of Protein Side Chain and Backbone Parameters from ff99SB. *Journal of Chemical Theory and Computation* **11**: 3696-3713.

Marvig, R.L., Johansen, H.K., Molin, S., and Jelsbak, L. (2013) Genome analysis of a transmissible lineage of *Pseudomonas aeruginosa* reveals pathoadaptive mutations and distinct evolutionary paths of hypermutators. *PLOS Genetics* **9**: e1003741.

Miller, V.L., and Mekalanos, J.J. (1988) A novel suicide vector and its use in construction of insertion mutations: osmoregulation of outer membrane proteins and virulence determinants in *Vibrio cholerae* requires toxR. *Journal of Bacteriology* **170**: 2575-2583.

Morán-Barrio, J., Lisa, M.-N., Larrieux, N., Drusin, S.I., Viale, A.M., Moreno, D.M. et al. (2016) Crystal Structure of the Metallo-β-Lactamase GOB in the Periplasmic Dizinc Form Reveals an Unusual Metal Site. *Antimicrobial Agents and Chemotherapy* **60**: 6013-6022.

Oliver, A. (2020) Antibiotic Resistance and Pathogenicity of Bacterial Infections Group - IdISBa

In.

Powers, R.A., Caselli, E., Focia, P.J., Prati, F., and Shoichet, B.K. (2001) Structures of Ceftazidime and Its Transition-State Analogue in Complex with AmpC β-Lactamase: Implications for Resistance Mutations and Inhibitor Design. *Biochemistry* **40**: 9207-9214.

Rodríguez-Martínez, J.-M., Poirel, L., and Nordmann, P. (2009) Extended-Spectrum Cephalosporinases in *Pseudomonas aeruginosa*. *Antimicrobial Agents and Chemotherapy* **53**: 1766-1771.

Roe, D.R., and Cheatham, T.E. (2013) PTRAJ and CPPTRAJ: Software for Processing and Analysis of Molecular Dynamics Trajectory Data. *Journal of Chemical Theory and Computation* **9**: 3084-3095.

Ryckaert, J.-P., Ciccotti, G., and Berendsen, H.J.C. (1977) Numerical integration of the cartesian equations of motion of a system with constraints: molecular dynamics of n-alkanes. *Journal of Computational Physics* **23**: 327-341.

Seabra, G.d.M., Walker, R.C., Elstner, M., Case, D.A., and Roitberg, A.E. (2007) Implementation of the SCC-DFTB Method for Hybrid QM/MM Simulations within the Amber Molecular Dynamics Package. *The Journal of Physical Chemistry A* **111**: 5655-5664.

Simon, R., Priefer, U., and Pühler, A. (1983) A Broad Host Range Mobilization System for In Vivo Genetic Engineering: Transposon Mutagenesis in Gram Negative Bacteria. *Bio/Technology* **1**: 784-791.

Tomalka, A.G., Zmina, S.E., Stopford, C.M., and Rietsch, A. (2013) Dimerization of the *Pseudomonas aeruginosa* translocator chaperone PcrH is required for stability, not function. *Journal of bacteriology* **195**: 4836-4843.
